# Experimental Tests of the Virtual Circular Genome Model for Non-enzymatic RNA Replication

**DOI:** 10.1101/2023.01.16.524303

**Authors:** Dian Ding, Lijun Zhou, Shriyaa Mittal, Jack W. Szostak

**Affiliations:** Department of Chemistry and Chemical Biology, Harvard University, 12 Oxford Street, Cambridge, Massachusetts 02138, USA; Department of Molecular Biology and Center for Computational and Integrative Biology, Massachusetts General Hospital, 185 Cambridge Street, Boston, Massachusetts 02114, USA; Department of Biochemistry and Biophysics, Perelman School of Medicine, University of Pennsylvania, Philadelphia, PA 19104, USA; Department of Genetics, Harvard Medical School, 77 Avenue Louis Pasteur, Boston, Massachusetts 02115, USA; Howard Hughes Medical Institute, Department of Chemistry, The University of Chicago, Chicago, Illinois 60637, USA

## Abstract

The virtual circular genome (VCG) model was proposed as a means of going beyond template copying to indefinite cycles of nonenzymatic RNA replication during the origin of life. In the VCG model the protocellular genome is a collection of short oligonucleotides that map to both strands of a virtual circular sequence. Replication is driven by templated nonenzymatic primer extension on a subset of kinetically trapped partially base-paired configurations, followed by shuffling of these configurations to enable continued oligonucleotide elongation. Here we describe initial experimental studies of the feasibility of the VCG model for replication. We designed a small 12-nucleotide model VCG and synthesized all 247 oligonucleotides of length 2 to 12 corresponding to this genome. We experimentally monitored the fate of individual labeled primers in the pool of VCG oligonucleotides following the addition of activated nucleotides, and investigated factors such as oligonucleotide length, concentration, composition, and temperature on the extent of primer extension. We observe a surprisingly prolonged equilibration process in the VCG system that enables a considerable extent of reaction. We find that environmental fluctuations would be essential for continuous templated extension of the entire VCG system, since the shortest oligonucleotides can only bind to templates at low temperatures, while the longest oligonucleotides require high temperature spikes to escape from inactive configurations. Finally, we demonstrate that primer extension is significantly enhanced when the mix of VCG oligonucleotides is pre-activated. We discuss the necessity of ongoing *in-situ* activation chemistry for continuous and accurate VCG replication.

## Introduction

Nonenzymatic RNA replication is thought to have been an essential early step that allowed the first RNA world protocells to begin the process of Darwinian evolution. As an intermediate stage between untemplated nucleotide polymerization and ribozyme-catalyzed RNA replication, template-directed nonenzymatic replication could have enabled the replication of protocells seeded with initially random sequences. Such a chemically driven exploration of sequence space would have set the stage for the evolution of the first functional ribozymes.^1^ Although recent advances have suggested potential routes for extensive template copying by RNA primer extension, going beyond template copying to cycles of replication remains a significant challenge.^2^

The nonenzymatic copying of a long RNA strand would result in a stable duplex that must be dissociated to allow for the next round of replication. A variety of potential solutions to the strand separation problem have been suggested. Thermal denaturation can only separate the strands of duplexes <30 bp under prebiotically plausible conditions.^1^ Other environmental influences such as pH fluctuations^3^ and microscale water evaporation/condensation cycles^4^ can potentially couple with thermocycling to facilitate strand separation. However, the reannealing of the separated strands is much faster than the rate of template copying at reasonable concentrations, which would block primer extension. Small fractions of backbone 2′-5′ linkages or DNA were shown to lower the melting temperature,^5–7^ but they also increase the hydrolytic lability of the duplex and slow down primer extension.^8,9^ As an alternative strategy, our lab has previously demonstrated that RNA oligonucleotides can lead to toehold mediated branch migration that can open up a segment of the duplex, allowing for strand displacement synthesis by nonenzymatic primer extension.^10^ This approach is closer to the helicase catalyzed strand displacement that occurs at replication forks in modern biology, but other problems with nonenzymatic RNA replication remain.

The difficulties in replicating ribozyme-length sequences recently led us to consider the assembly of functional ribozymes by the ligation of shorter oligonucleotides that would be easier to replicate. Our lab has recently demonstrated that splinted ligation and loop-closing ligation can form functional ribozymes from short oligonucleotides.^11,12^ However, even the replication of shorter oligonucleotides faces problems that can lead to information loss at both ends of the sequence. First, the nonenzymatic copying of the last base of a sequence by primer extension is known to be very slow relative to the copying of internal nucleotides,^13^ which could lead to progressive loss of 3ʹ-sequences over cycles of replication. This notorious “last base addition problem” is now understood as being due to the primary mechanism of nonenzymatic primer extension, which requires the binding of an imidazolium-bridged dinucleotide intermediate (N*N) to the template by two base-pairs.^14^ With only one base pair possible at the last base of the template, binding of the bridged dinucleotide is greatly weakened, thus reducing the rate of primer extension. While an imidazole-activated mononucleotide can still perform nonenzymatic primer extension, the reaction is much slower and more error-prone.^2,15^

Maintenance of the genetic information at the 5′-end of a sequence is even more problematic, since this would require a continuous supply of a specific primer, which is clearly not prebiotically plausible. As a result, information will be lost when nonenzymatic primer extension is initiated at an internal position on a template. Although ligation events could potentially salvage some internally initiated strands, this process is slow and inefficient, and would be completely prevented if the 5′-end is unphosphorylated or is blocked by a nucleotide 5′-5′-pyrophosphate cap.

These problems have led others to propose that primordial genome replication occurred by a rolling circle process, in which primer extension continues many times around a circular template, spinning off a long multimeric single stranded product.^16,17^ As in modern viroid replication this linear product would have to be cleaved into unit length strands, which would then have to become circularized to generate a circular template. The process would then have to repeat for the other strand. Since this process would require very extensive primer extension in the face of the topological difficulties of replicating a small circular RNA, as well as requiring multiple ribozyme activities for cleavage and circularization, we do not consider rolling circle replication to be a viable model for nonenzymatic RNA replication.

The above problems led us to propose the Virtual Circular Genome (VCG) model for prebiotically plausible nonenzymatic RNA replication.^18^ As described in our VCG hypothesis paper, the primordial genome is comprised of a collection of short oligonucleotides that map onto both strands of one or more virtual circular sequences (Figure 1A). Since a circular genome does not have a defined start or end, copying can be initiated and terminated at any position. This genome is not represented by any actual circular molecules, but is instead represented by all possible fragments from both strands of the virtual sequence. In theory, every oligonucleotide in this system can act as a primer, template, or as a downstream helper due to stacking interactions or by forming an imidazolium-bridged intermediate. Denaturation and reannealing induced by environmental fluctuations can generate kinetically trapped partially basepaired configurations,^19^ of which a productive fraction will enable primer extension and ligation to occur (Figure 1B). Shuffling of these base-paired configurations would allow for additional elongation to occur, and RNA mediated branch migration could also open up base-paired regions, allowing for primer extension by strand displacement synthesis. In this model, the process of genetic replication is distributed across all of the oligonucleotides of the entire system through cycles of rearrangements of base-paired configurations.

**Figure 1.**
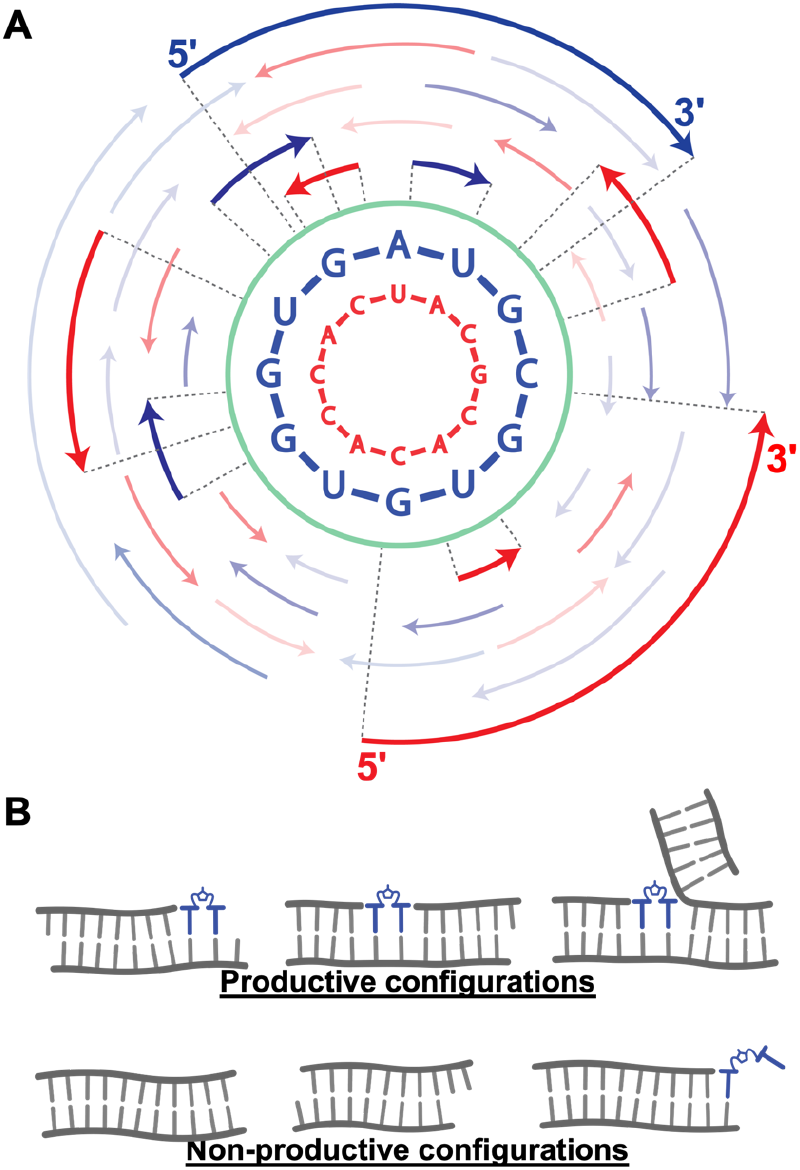
(A) Schematic illustration of the virtual circular genome model. The green circle represents the virtual genome that does not correspond to any actual oligonucleotide. A subset of the real oligonucleotides in the VCG system is illustrated as the blue and red arrows. Dotted lines along with the bold arrows showed how the oligomers map onto the virtual circular genome. The two complementary sequences selected for this study is shown inside the green circle. The direction from 5ʹ to 3ʹ is clockwise for the blue sequence and counterclockwise for the red sequence, which is the same direction as the arrows. (B) Examples of productive and non-productive configurations of the oligonucleotides arrangement.

We envision the VCG system as in effect an assembly line where newly generated or introduced short oligonucleotides gradually become elongated to strands of roughly 10-20 nucleotides in length. Oligonucleotides of this length can then be assembled into functional ribozymes, either by splinted ligation^11^ or by iterated loop-closing ligation.^12^ These ribozyme building blocks could be the end products of one or potentially multiple virtual circular genomes replicating together in a protocell in a prebiotically plausible environment.

Here we explore a model VCG system with a 12-nt long virtual genome represented by 247 different oligonucleotides which range from 2 to 12 nucleotides in length. Several dimers and trimers occur multiple times in the sequence. Using radiolabeling, we monitored the fate of individual oligonucleotides in the system following the addition of activated nucleotides or bridged dinucleotides. We investigated the effect of factors including oligonucleotide length, concentration, and temperature on the primer extension yield. In the course of these studies, we discovered a surprisingly prolonged equilibration process of the oligonucleotide mix in the VCG system that enables a considerable extent of reaction. Furthermore, we found that temperature fluctuations would be essential for continuous and templated extension of the entire VCG system across different oligo lengths. Finally, we discuss the necessity of either a flow system or ongoing *in-situ* activation chemistry for continuous and accurate VCG replication.

## RESULTS

### Primer extension in the VCG mix versus on a single template

To begin to test the virtual circular genome model, we first selected a 12-nt virtual circular genome sequence with no secondary structure or kinetically severe stalling points such as UU sequences that are difficult to copy (Figure 1A). The sequence that we selected is represented by 247 different oligonucleotides, ranging from 2 to 12 nucleotides in length, that map to either strand of the virtual circular sequence (Table S1). Every oligonucleotide in the system can in principle bind to many complementary oligonucleotides, but the most thermodynamically favored pairing will be the formation of a fully base-paired duplex. In order to form kinetically trapped partially base-paired configurations for template copying, we used a brief (10 s) initial 90°C pulse to disrupt all base-pairing. We expected subsequent fast cooling to trap a fraction of the oligonucleotides in metastable configurations that would allow complementary imidazolium-bridged dinucleotides to bind to a template strand next to a primer and react by primer extension (Figure 1B). Imidazolium-bridged dinucleotides can extend a primer by one nucleotide, with an activated mononucleotide displaced as the leaving group. All ten possible intermediates were supplied at the same concentration (∼1.7 mM each) for all primer extension reactions in the system (Figure S1).

We then set out to determine whether it is possible for oligonucleotides to be elongated by primer extension in the highly complex virtual circular genome system. We monitored the extension of individual labeled primers occurring within the mixture of 247 different VCG oligonucleotides (Figure 2A). We started by monitoring a single radiolabeled 6-mer oligonucleotide added in trace concentration (< 0.05 μM) to a mixture of 1 μM of each VCG oligonucleotide, which we refer to as the 1X VCG mixture. About half of the initial radiolabeled 6-mer was extended to the corresponding 7-mer in 1 day. This rate of primer extension was much slower than in the positive control in which the same labeled primer was incubated with only one complementary 12-mer template. Nevertheless, this observation shows that a significant fraction of the VCG oligonucleotides anneal to form configurations that are productive for primer extension, and that a fraction of these configurations exist for a time scale of hours to days.

**Figure 2.**
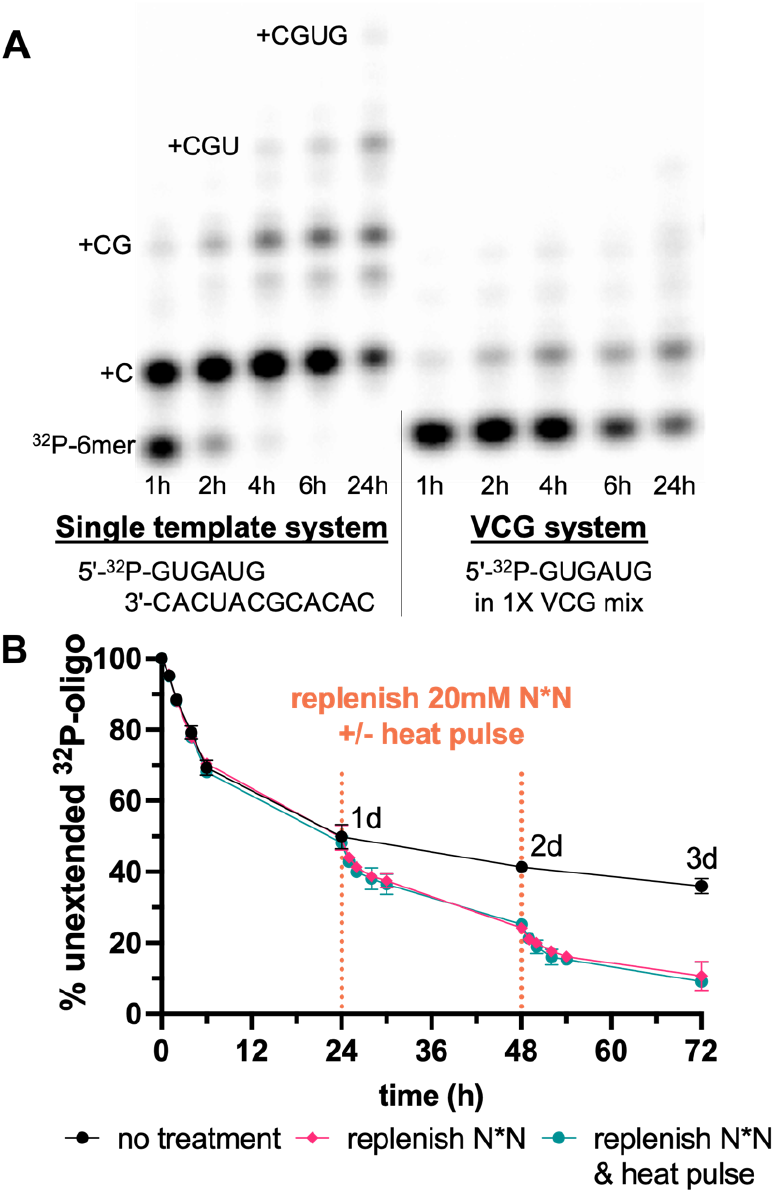
Demonstration of extension inside the virtual circular genome system by a labeled 6-mer: 5ʹ-^32^P-GUGAUG. (A) Comparison between the VCG system and a single template system. (B) Continuous VCG extension in 3 days, with and without further replenishment of activated N*N and 90°C heat pulses. The VCG system contains 1 μM of all the VCG oligos listed in Table S1. The single template system contains 1μM of template and 1 μM of primer, dopped with 5ʹ-^32^P-GUGAUG. All reactions were conducted at room temperature, with 50 mM MgCl_2_, 200 mM Tris-HCl (pH 8.0), and 20 mM pre-equilibrated N*N.

We then asked what limits primer extension in the virtual circular genome system compared to the single template system. One possibility is that fast equilibration of the oligonucleotides depletes available templates as they become sequestered in stable duplexes. Since only one heat pulse was applied to dissociate duplexes and initialize the process, if all oligonucleotides with melting points above room temperature quickly equilibrated back to form stable duplexes with their own complementary strands, then the labeled 6-mer would be rapidly disassociated from any suitable templates for primer extension. An alternative extreme possibility would be continued but very slow rearrangement of the initially formed oligonucleotide complexes. If all oligonucleotide configurations after the initial heat pulse were locked in place, then any radiolabeled 6-mer trapped in an unproductive configuration would not be able to shuffle into a productive configuration, and primer extension would cease after all initially productive configurations had become extended. However, it is unlikely for a 6-mer with an estimated k_off_ of ∼19 s^-1^ to its complementary strand^19^ to bind so tightly that it could not either spontaneously dissociate from its template or be strand displaced by another longer complementary oligonucleotide. The resulting free 6-mer could then anneal to a new template, where it would have another opportunity to be extended. Besides the equilibration and rearrangement rates, another potential limiting factor in the VCG system is simply the proportion of productive configurations at any given time. Since nonenzymatic templated extension requires at least two open nucleotide binding sites downstream of a template-bound primer for efficient reaction, any other kinetically trapped configurations will block templated extension (Figure 1B). Unlike the single template system, where most primers can form the appropriate primer-template complex and therefore be extended, many of the oligonucleotides in the VCG system will be at least initially bound in unproductive configurations. Because the initial rate of primer extension in the VCG mix is slower than the rate in the single-template control, we hypothesize that the initial limiting factor for fast primer extension is the proportion of productive configurations, and that slow equilibration in the complex virtual circular genome system as well as the ongoing hydrolysis of the activated species are responsible for the subsequent continuing decline in the rate of primer extension.

### Rearrangement and equilibration of the base-paired configurations in VCG system

To test the idea that continued spontaneous shuffling of productive configurations might be occurring, we allowed the same primer extension reaction to continue for an extended time without any external treatments. Remarkably, template directed primer extension continued for at least three days, at an ever-declining rate. (Figure 2B) This result suggests that at least a fraction of the oligonucleotide complexes was still shuffling and acting as templates for primer extension after three days. However, we suspected that the declining primer extension rate was also partially due to a declining concentration of activated species (N*N bridged dinucleotides) available at later times because of their relatively rapid hydrolysis under primer extension conditions (t_½_ ≈ 5 h) (Figure S2). Therefore, we performed a similar 3-day VCG reaction with replenishment of N*Ns each day. These freshly supplied activated species boosted the extent of primer extension in the VCG system, suggesting that a significant proportion of productive oligonucleotide configurations were still present in the system after three days. To our surprise, when additional heat pulses were performed just prior to each N*N replenishment, no significant improvement in primer extension was observed. We speculate that the medium-sized oligonucleotides in the VCG system were probably shuffling well enough at room temperature to continuously generate productive configurations that additional heat pulses to reset the system did not induce significant improvement.

Given the remarkably prolonged equilibration process in the VCG model, we asked if system-wide changes in oligonucleotide concentrations would impact the observed extent and rate of primer extension. Diluting or concentrating the entire VCG oligo mixture will affect the concentration of every oligonucleotide complex in the system by affecting the association rate for duplex formation. Although one might expect that dilution, and hence weaker binding of the short 6mer primer to templates would result in reduced primer extension, what we observed was the opposite. Under the same reaction conditions, a less concentrated VCG mixture exhibited faster primer extension and a greater yield of extended product (Figure 3A). We suggest that the lowered concentration of short oligonucleotides allowed for a greater initial fraction of productive configurations, and that the slower association rate for duplex formation allowed newly opened templates to remain available for primer extension for a longer time. This result suggests that concentration fluctuations could facilitate the continued rearrangement of oligonucleotide configurations in the VCG mix. Changes in oligonucleotide concentration can also be interpreted in terms of concentration dependent changes in duplex T_m_. A more dilute VCG mix implies a lower effective T_m_ for all oligonucleotide duplexes, which could facilitate continued shuffling of base-paired configurations. As a point of reference, we measured the melting temperature of our 6-mer primer and its complement at three different concentrations in primer extension buffer to demonstrate this relationship (Figure 3A). A 3-fold decrease in concentration led to a 1°C decrease in T_m_, and even this modest effect was enough to lead to a noticeable increase in primer extension.

**Figure 3.**
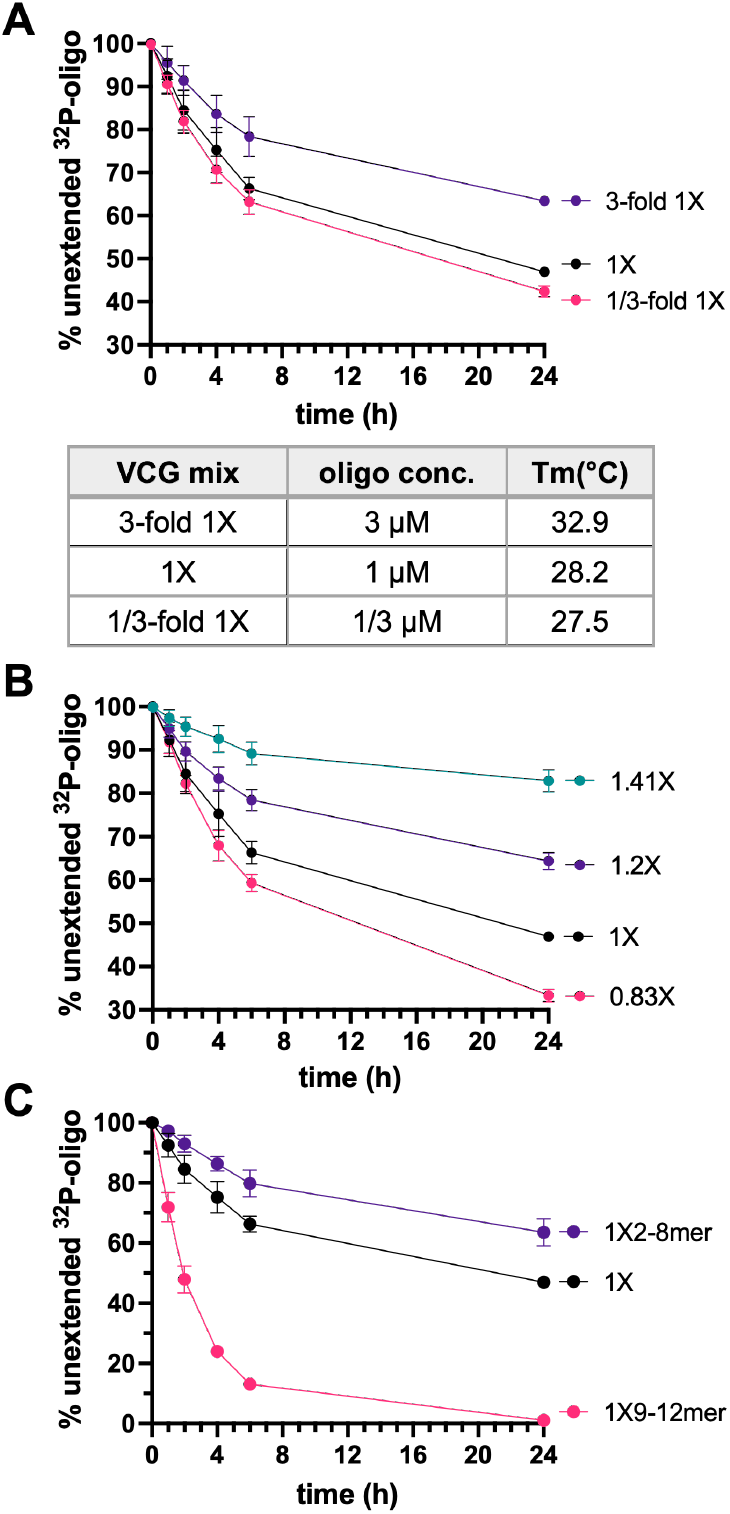
Virtual circular genome extension with different oligomers compositions. (A) VCG extension when concentrated and diluted. The bottom table indicates the concentration of each oligomer in these three mixtures and the melting temperature of pGUGAUG measured at the indicated concentration. (B) VCG extension at different concentration gradient. The concentration gradient is expressed as [(pN)_i_]/[(pN)_i+1_], starting at 1 μM of each 12mer. (C) Extension in partial VCG mixtures containing only the longer or shorter oligomers. All reactions were measured by the extension of 5ʹ-^32^P-GUGAUG (< 0.05 μM) conducted at room temperature, with 50 mM MgCl_2_, 200 mM Tris-HCl (pH 8.0), and 20 mM pre-equilibrated N*N. See Table S2 for detailed oligomer concentrations in different VCG mixtures.

In a further attempt to manipulate the proportion of productive configurations in the VCG system, we adjusted the concentrations of the VCG oligonucleotides in a length dependent manner. We reasoned that if on average elongation by primer extension is slow, as we observed, then a length dependent concentration gradient might emerge, with shorter oligonucleotides being more abundant than longer oligonucleotides. For the following experiments we made the simplifying assumption of an exponential gradient of length distribution, where the concentration gradient is defined as [(pN)_i_]/[(pN)_i+1_]. For example, a 1.41X VCG system with 1 μM of each 12-mer contains 1.41 μM of each 11-mer and 2 μM of each 10-mer, etc. Table S2 lists the concentration of each oligonucleotide as a function of length in the different concentration gradients that we used for the following experiments. As previously noted, with a 2X concentration gradient, primer extension of all oligonucleotides by one nucleotide on average results in duplication of the entire population, i.e. one round of replication. Similarly, a 1.41X (≈√2) gradient requires an average of 2 nucleotides and a 1.2X (≈∜2) gradient requires approximately 4-nt of primer extension for one round of replication.^18^

Experimentally, we observed that a steeper concentration vs. length gradient leads to a significantly slower rate of extension of a labeled 6-mer primer (Figure 3B). We interpret this effect as being due to increased competition for binding to the limited concentration of longer oligonucleotides, which are expected to be better templates as they are long enough to provide binding sites for a primer, a bridged dinucleotide substrate and a downstream helper. The ratio of a 6-mer primer to a 12-mer template in a 1X concentration gradient is 1:1, but this ratio increases to 8:1 in the 1.41X and 64:1 in a 2X concentration gradient. As a result, the fraction of the 6-mer primer that is able to bind to a longer template oligonucleotide is lower with a steeper gradient. Thus primer extension, expressed as a fraction of input primer, is decreased; however it should be noted that the total amount of extended primer is increased. For example, while the 1X gradient can produce 1 μM × 53% = 0.53 μM of newly extended 7-mer in one day, the 1.41X gradient can produce up to 8 μM × 18% = 1.44 μM, almost triple the amount. The effect of concentration gradient on the extension rate is seen with oligonucleotides of different lengths. We measured the extension of 8-, 10- and 12-nt primers, and in all cases the fraction of primer extended vs. time was higher in a VCG mix with a shallower concentration vs. length gradient (Figure S3). We also tested a 0.83X gradient, where longer oligonucleotides are present at higher concentrations than shorter oligonucleotides. With this reverse gradient, we observed a faster rate of primer extension than in a 1X gradient, presumably due to the higher availability of longer oligonucleotides as good templates.

To further investigate the factors controlling the rearrangement of base-paired configurations, we explored partial VCG systems where only the shorter or longer VCG oligonucleotides were supplied. An optimal template for primer extension requires sufficient complementarity to the primer for stable binding, and at least two additional unpaired nucleotides downstream of the primer to act as the binding site for an activated bridged dinucleotide. For our 6-mer primer, an optimal template would need to be at least 8-nt long. We first examined a partial VCG system consisting of only 2-to 8-mer oligonucleotides. In this system, only one of the 24 8-mers would be an optimal template for the radiolabeled 6-mer. We observed a slower initial rate of primer extension, and a lower extent of primer extension at 24 h in the 2-8mer partial VCG system than in the complete system (∼36 % vs. 53 %), presumably because of the low proportion of productively arranged 6-mer primer at any given time point. (Figure 3B) However, even though the rate was low, this observation suggests that even a VCG system with an 8-nt genome allows extensions. On the other hand, the 9-12mer partial VCG system which contains only the longer subset of oligonucleotides shows extremely good primer extension, with essentially complete primer extension by one or more nucleotides in one day. Because all of these longer oligonucleotides are present together with their complementary strands in the VCG system, we initially expected that the rapid formation of stable duplexes would prevent significant primer extension. Since not all of the radiolabeled 6-mer could be in a productive configuration after the initial heat pulse, the fact that primer extension continued until all of the 6-mer primer had been extended implies that rearrangements of base-paired configurations were happening in the VCG system for these 9-12nt oligonucleotides at room temperature.

### Extension of oligonucleotides of different lengths in the VCG system

The length of an oligonucleotide in the VCG system is likely to affect both its initial likelihood of annealing in a productive configuration, as well as the dynamics of the exchange processes that would allow for continued primer extension. We therefore determined primer extension rates for a series of oligonucleotides of different lengths (Figure 4A). In order to avoid effects of differing sequences at the 3′-end of the primer, we used a set of oligonucleotides with the same 3′-end as the 6-mer primer used above, and varied only the 5′-end. Initially we expected that longer oligonucleotides might show faster initial rates of primer extension, since they would be able to bind more strongly to longer templates. We also expected slower long-term rates of primer extension since they would be more likely to become sequestered in stable unproductive configurations that would be unable to exchange into new productive configurations. Surprisingly, we observed a progressive decrease in both the initial and long-term rates of primer extension as oligonucleotide length increased from 6 to 8, 10, and then to 12 nucleotides. We suggest that both of these effects stem from a decreased probability of forming productive configurations. The melting temperature of these oligonucleotides when paired with their perfect complement increases significantly with length (Figure 4B). This greater duplex stability is likely to decrease spontaneous shuffling of paired configurations in the VCG system, decreasing the rate of primer extension at long times. Why longer primers are extended more poorly initially is less clear, but could potentially be due to occupancy by pairs of shorter oligonucleotides preventing the formation of productive configurations. Alternatively, toehold mediated branch migration may lead to rapid loss of productive configurations, thereby decreasing primer extension even at early times.

**Figure 4.**
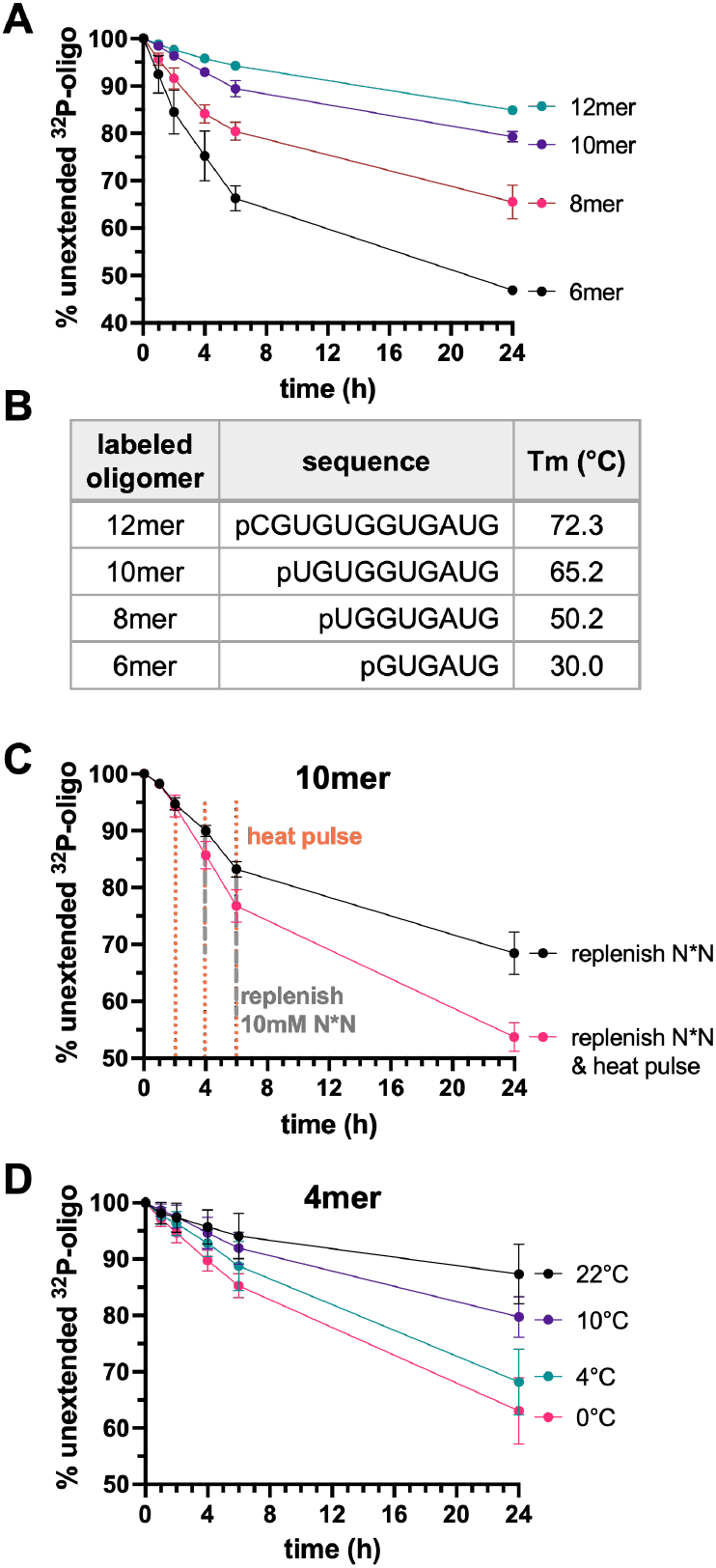
Length dependency and temperature effect of extension inside VCG system (A) Extension of oligomers with different length in 1X VCG oligo mix, represented by the percentage of unextended 5ʹ-^32^P-labeled oligonucleotide over time. (B) Sequences of the labeled oligomers and their melting temperatures, measured at primer extension buffer. (C) Heat pulses facilitate the continuous VCG extension of a 10mer (5ʹ-^32^P-UGUGGUGAUG). The sample with heat pulses were heated up to 90°C for 10 s and immediately cooled on ice for 1 min at 2, 4, 6 h. The two experiments were replenished with 10 mM of equilibrated and lyophilized N*N at 4 h and 6 h. (D) Lower temperature facilitates the VCG extension of a 4-mer (5ʹ-^32^P-GAUG). All reactions were conducted at room temperature, 1X VCG oligo mix, 50 mM MgCl_2_, 200 mM Tris-HCl (pH 8.0), and started with 20 mM pre-equilibrated N*N.

To facilitate shuffling of the longer oligonucleotides for continuous elongation, we tested the effect of periodic temperature fluctuations on extension of the 10-mer primer. After an initial high temperature pulse to initialize the system, three additional high temperature pulses (90 °C for 10 s) were applied every two hours to shuffle the oligonucleotide configurations. Fresh activated N*Ns were added after the second and third high temperature pulse to counter the effects of hydrolysis, since about half of the initial bridged dinucleotides had already hydrolyzed after 4h (Figure S1-2). A clear improvement in the extent of 10-mer extension was observed with the extra high temperature pulses, demonstrating the importance of temperature fluctuations for the continued elongation of longer oligonucleotides in the VCG system (Figure 4C). As expected, the improvement in primer extension was even greater when fresh N*N substrates were added after each high temperature pulse.

In contrast to the requirement of high-temperature fluctuations for primer extension of longer oligonucleotides, we have found that shorter oligonucleotides can only be extended at lower temperatures. When a 4-mer primer is radiolabeled and monitored in the VCG mix at room temperature (22°C), we observe only minimal primer extension. In addition to the low yield, many of the extended products formed were incorrect (Figure S4B). We first hypothesized that in the VCG mix, much of the 4-mer was not bound to any template most of the time at room temperature, and that the observed extension arose primarily through non-templated extension. However, a control experiment showed that the 4-mer can extend efficiently on a single template at room temperature (80% at 24 h), although the extent of primer extension does improve markedly at lower temperatures (Figure S4A). Therefore, the poor extension of the 4-mer primer in the VCG system is not solely due to poor binding. We speculated that rapid disassociation of the 4-mer from a template strand, followed by template occupancy by a competing oligonucleotide, would prevent primer extension. We therefore tested the effect of reducing the temperature on 4-mer extension yield in the VCG system. As decreased temperatures, we observed increased correct extension and decreased misincorporation (Figures 4D, S4B). The remarkably improved yield and fidelity suggests that primer extension of the shorter oligonucleotides in the VCG system requires a lower temperature to prevent rapid loss of productive configurations.

### Fidelity in the virtual circular genome scenario

The significant degree of misincorporation observed with the 4-mer at room temperature drew our attention to the possibility of untemplated extension in the VCG system. Untemplated extension could result not only from unbound oligonucleotides but also from some of the unproductive configurations in the VCG system. As shown in figure 1B, many unproductive configurations have either an overhanging or blunt 3′-end that can potentially be subject to untemplated extension. Moreover, the 5′-phosphate of both free oligonucleotides and some template-bound oligonucleotides may also react to form 5′-5′-pyrophosphates. Indeed, PAGE analysis of extension of the 6-mer primer in the VCG system (Figure 2A) clearly shows that several products are formed that are not seen in the single template system, suggesting that these misincorporations most likely derive from processes other than templated primer extension. Interestingly, increasing the concentration of the bridged dinucleotides enhanced the synthesis of these incorrect products, but did not significantly improve the correct templated primer extension reaction (Figure S5).

In order to identify the sources of these misincorporations, we first examined untemplated extension of specific oligonucleotides in the presence of all possible activated bridged dinucleotides. In the VCG system, because every oligonucleotide exists in the presence of their partially and fully complementary strands, blunt ends can form at either end of the oligonucleotide. Therefore, we tested both single-stranded RNAs of different lengths and the corresponding double-stranded duplexes for untemplated extension. To our surprise, we observed enhanced untemplated extension with blunt ended species (Figure S6).

Because of the limited ability of PAGE analysis to resolve different products of untemplated extension, we determined the extent and regioselectivity of untemplated extension by suppling only one bridged homo-dinucleotide at a time (Figure S7). The identity of each extended product was determined by comparison with authentic radiolabeled samples (see Materials and Methods for synthesis of standards). Untemplated oligonucleotide polymerization has long been known to favor 2′-over 3′-extension due to the greater nucleophilicity of the 2′-hydroxyl group, and the formation of 5′-5′ pyrophosphate products is known to be an unavoidable byproduct of reactions with nucleotide phosphorimidazolides.^20,21^ In our examination of untemplated extension, we also observed a predominance of products with nucleotides added at either the 2′-OH or the 5′-phosphate. Blunt ended duplex oligonucleotides appear to be particularly prone to nucleotide addition to the 2′-hydroxyl, especially with G (Figure S7). In addition to the untemplated extension of single-stranded and blunt end RNAs, we also examined primer extension of the labeled 6-mer primer in VCG mix in the presence of only one imidazolium-bridged homo-dinucleotide at a time. Note that correct templated extension in this case requires a C*G bridged dinucleotide. In the absence of this fully complementary substrate, the products of primer extension were quite similar to those of the untemplated reactions, with most of the elongations being at the 2′-or 5′-end. When supplied with a C*C bridged dinucleotide the observed correct 3′-extension with C probably resulted from the substrate binding to the template with a downstream C:C mismatch. Interestingly, we observed less extension with bridged homo-dinucleotides in the VCG system than with an isolated duplex, especially when G*G was supplied. This finding suggests that annealing of the oligonucleotides in the VCG mix results in a low proportion of blunt ended duplexes, as might be expected since there are many more annealed configurations with 5′-or 3′-overhangs than blunt ended configurations.

To better identify the misincorporation, we aligned the gel separated VCG extension products with the individual untemplated reaction products; we also used phosphatase digestion to distinguish between the 5ʹ-(which protects the ^32^P-labeled 5′-phosphate from digestion) and 2ʹ/3ʹ-extension (Figure 5). This assay showed that most of the apparent misincorporations in the VCG reaction were in fact due to 5′-nucleotide-pyrophosphate formation. We could not quantify how much of each pyrophosphate is formed because primers with a 5′-App-, 5′-Upp and 5′-Cpphave almost identical gel mobilities. However, in the reactions with single N*N substrates, A*A led to more formation of 5′-App-oligo products than the corresponding products with C*C, U*U and G*G. It is also possible that 5′-Gpp extension of the specific radiolabeled primer we used could be template directed in the VCG mix. The 2′+A and 2′+U products have similar gel mobility to the correct (templated) 3′+C product. However, we believe that there is little 2′-extension with A and U because no significant amount of 2′+C or 2′+G products were formed (Figure 5&S7B). The small amount of slowly migrating products in the VCG prime extension reaction that survived the phosphatase digestion likely correspond to the 5′-Npp extension of the correct 3′+C product. As a result, most of the misincorporations we observed in the VCG systems appear to result from 5′-5′-pyrophosphate formation. We note that 5′-Npp-capped oligonucleotides can still act as fully functional primers and templates in the VCG system; the accumulation of 5′-Npp-oligonucleotides could also provide a selective advantage for the evolution of ribozyme ligases that use such molecules as susbrates.^22^

**Figure 5.**
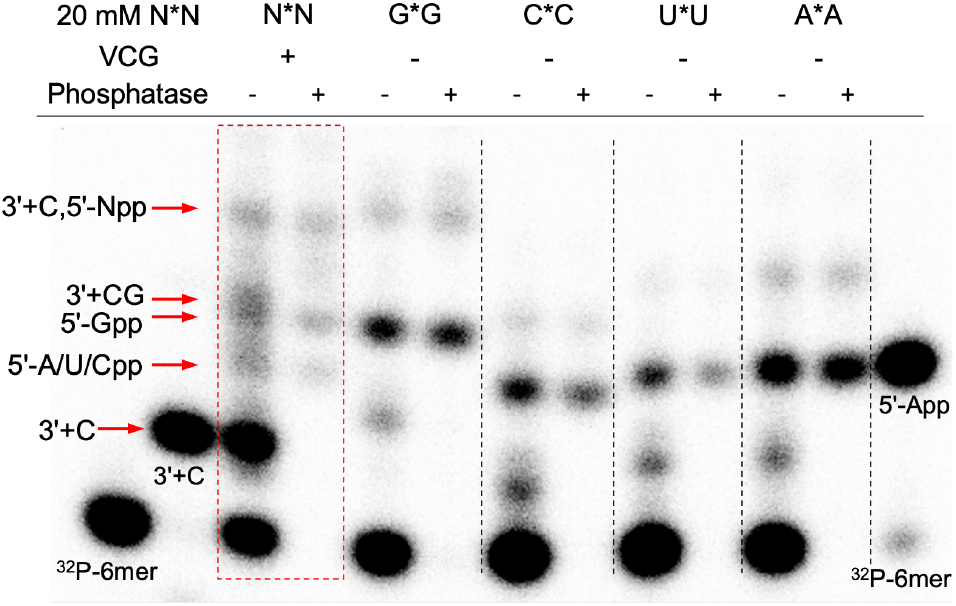
PAGE gel analysis of the products from phosphatase digestion. The 5ʹ-^32^P-oligonucleotides were de-radiolabeled while the 5ʹ-Np^32^P-oligonucleotides were protected. The VCG extension were performed with 1X VCG mixture and 20 mM N*N, while the nontemplated reactions were ran with 1 μM 6mer and 20 mM of the indicated imidazolium-bridged homodimers. All reactions were run at room temperature for 24h. Phosphatase digested products were loaded at the same concentration as the untreated sample. Authentic samples were run alongside on the PAGE gel and indicated in the figure.

### Potential strategies to enhance extensions in a VCG system

Having characterized the basic kinetics and fidelity of primer extension in our model VCG system, we ask what factors might further increase the rate and yield of primer extension. Considering the rapid hydrolysis of imidazolium-bridged substrates, an efficient method for *in-situ* activation would likely be extremely beneficial. This ideal approach would lead to efficient activation of both monomers and oligonucleotides, as this would allow for the formation of monomers bridged to oligonucleotides, which we previously shown to be optimal substrates for primer extension.^2^ We therefore asked whether pre-activation of the VCG oligonucleotide mix would enhance primer extension by allowing for the formation of monomer-bridged-oligonucleotide intermediates.

We began by testing whether an activated trimer helper can accelerate the extension of our labeled 6-mer primer in the VCG system. We prepared the activated trimer *GUG and doped it at increasing concentrations into the partial 9-12mer VCG system. Following the addition of activated monomers or bridged dinucleotides, this trimer can form the highly reactive C*GUG intermediate *in-situ*. The higher affinity and greater pre-organization of this substrate facilitates the +C extension of the ^32^P-labeled 6-mer primer. Previous kinetic measurements have shown that a similar monomer-bridged-trimer (specifically, A*CGC) has a K_M_ of 40 μM, and a Vmax approaching 1 min^-1^ on a single template.^2^ When we added the *GUG helper together with an equilibrated mix of imidazolium-bridged dinucleotides to the partial 9-12mer VCG oligonucleotides, we observed significant acceleration of primer extension when it was present at a concentration (∼50 μM) closer to the estimated K_d_ of C*GUG. Moreover, primer extension in the partial VCG system supplied with 100 μM *GUG can be almost as fast as the one-template positive control (Figure S8).

Encouraged by the observed benefit of adding a single activated helper oligonucleotide, we asked whether activating the entire set of VCG oligonucleotides would also help monomer-bridged-oligos to form *in-situ* and thus enhance primer extension. An important concern is that the excess amount of 2-aminoimidazole required for efficient activation will also reduce the formation of imidazolium-bridged substrates. To avoid this problem, we used stochiometric 2AI to activate a concentrated VCG oligonucleotide mix using partial freezing for eutectic phase and then dilute that for extension measurement. Eutectic phase concentration, followed by dilution for primer extension measurements. Partially freezing lead solute concentration in the liquid eutectic phase; we have previously used this approach to enable efficient in-situ activation of imidazolium bridged species for nonenzymatic template copying. Here we used the non-prebiotic 1-ethyl-3-(3-dimethylaminopropyl)carbodiimide (EDC) as the coupling reagent for activation for ease of handling, but similar activation chemistry can be performed using the more prebiotically plausible methyl isocyanide. A control NMR experiment with a single dinucleotide demonstrated almost complete activation under the same conditions (Figure S9). After overnight eutectic phase activation, the reaction was warmed to room temperature, diluted into primer extension buffer, and ^32^P-labeled 6-mer primer was added. We started by activating the 1.41X VCG mix, in which short oligonucleotides are present at higher concentrations than the longer oligonucleotides. We observed significant enhancement of primer extension (Figure 6A) even though the concentration of the short oligonucleotides were still far below the K_d_ of the corresponding monomer-bridged-oligomers.^2^ When we activated the 1X VCG system we observed no rate enhancement, probably because the concentration of the short helper oligonucleotides was too low. We then reasoned that the optimal concentration vs. length distribution might be more complex than a simple linear gradient. Clearly, short oligonucleotides must be present at close to their K_d_ to have a significant effect on prime extension. On the other hand, medium length oligonucleotides are elongated most rapidly, and therefore might be present at lower steady state concentrations, while longer oligonucleotides might accumulate and reach higher concentration. We therefore prepared and activated a VCG mix with a U-shaped concentration vs. length distribution (Table S2). We were pleased to observe improved primer extension in this system, with about 70% of the labeled primer being extended by one or more nucleotides in less than one day (Figure 6B).

**Figure 6.**
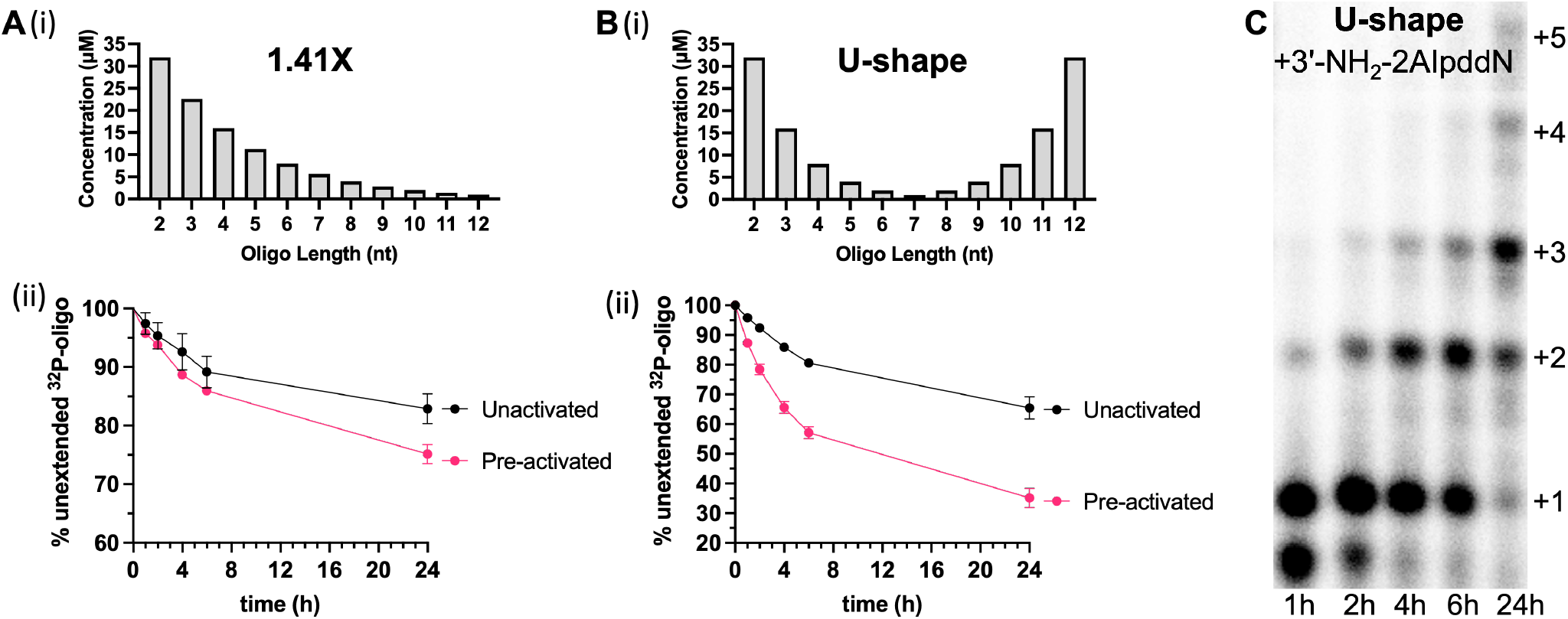
Demonstration of possible strategies to improve VCG extension (A-B) Significantly enhanced VCG extension after pre-activation with either 1.41X or U-shape gradient. (i) Oligo concentration of each length (ii) Comparison between the extensions of the 5ʹ-^32^P-GUGAUG inside the VCG system with or without the pre-activation (C) Faster extension in a U-shaped VCG mix with the more reactive 3ʹ-NH_2_-2AIpddN modification as a model system

Finally, we asked how primer extension in the VCG system would be affected if the reaction kinetics were improved. To do this we employed a ^32^P-labeled 6-mer primer terminated with a highly reactive 3′-amino-2′,3′-dideoxy-ribonucleotide, and similarly modified 2-aminoimidazole activated mononucleotides (3ʹ-NH_2_-2AIpddNs). Although such nucleotides may not be prebiotically plausible, they provide an excellent model system for the simulation of nonenzymatic RNA copying under conditions leading to enhanced rates of primer extension, such as might be achieved e.g. by a prebiotic catalyst or improved conditions for chemical RNA copying. By employing the highly nucleophilic 3′-amino group, we were able to observe ∼60% +1 primer extension in just one hour, with almost complete +1 or greater extension by 4 h, and a low fraction of misincorporations (Figure7C). Remarkably, average extension of ∼ +3 nucleotides was observed by 24 h, consistent with spontaneous shuffling of partially base-paired configurations continuing for many hours.

## DISCUSSION

We first proposed the virtual circular genome model^18^ as a theoretical means of overcoming the barriers to prebiotically plausible RNA replication. Replication in the VCG model does not require the specific primers needed for replication of a linear genome, and the distributed nature of the copying processes is expected to impart resilience to chemical processes that modify or block the 5′-or 3′-ends of individual oligonucleotides. Importantly, the repeated shuffling of base-paired configurations of annealed oligonucleotides was proposed as a means of overcoming the block to replication imposed by rapid strand annealing. However, experimental tests of this model were clearly needed, as template copying by primer extension has previously been examined only in highly simplified model systems. Our studies show that primers of different lengths can indeed be extended by template copying with a significant rate, extent and fidelity in a model VCG system, suggesting that under appropriate environmental conditions, replication in the VCG mode may be possible.

Perhaps the most surprising aspect of our results is the prolonged time scale (> 1 day) over which primer extension in the VCG mixture continues. We interpret the extended time scale of primer extension as reflecting the very slow equilibration of the VCG oligonucleotides. The very large number of competing base-paired configurations of VCG oligonucleotides may prevent rapid equilibration to fully base-paired duplexes, thus allowing for the continued shuffling of partially base-paired configurations. At any given time, only a fraction of these configurations is productive for primer extension, while others are not. If unproductive configurations can rearrange by dissociation, exchange or strand displacement, new productive configurations may continue to arise, enabling the observed extended time course of primer extension.

We have found that oligonucleotides that were both longer (8, 10 and 12-nts) and shorter (4-nts) than the 6-mer primer exhibited slower and less extensive primer extension. Short pulses of high temperature partially rescued the poor extension of the longer primers, suggesting that these oligonucleotides tend to become trapped in stable unproductive configurations that can be disrupted and exchanged during exposure to elevated temperatures. In contrast, the shorter 4-nt oligonucleotide required a lower temperature for optimal primer extension, in part due to the weaker binding to template strands, but also in part due to the greater lability of productive configurations involving a base-paired 4-nt primer. These divergent temperature requirements for primer extension of oligonucleotides of different lengths imply that repeated cycles of RNA replication would only be possible in a fluctuating environment. For example, changing temperature, pH or salt concentrations could trigger ongoing shuffling of the annealed configurations of the VCG oligonucleotides.

The observed variation in the rate of primer extension with primer length has implications for the steady state length vs. concentration profile. In our original model, we assumed for simplicity an exponential concentration vs. length gradient, i.e. a constant [length n]/[length n+1] ratio. An attractive feature of such simple models is that overall replication can be achieved through the elongation of all oligonucleotides in the population by only 1 or 2 nucleotides. However, if short and long oligonucleotides are elongated more slowly than medium-length oligonucleotides, the steady state length vs. concentration distribution may be more U-shaped, because both short and long oligonucleotides would tend to accumulate, while medium-length oligonucleotides would be rapidly extended and converted into long oligonucleotides, which would then accumulate. The short oligonucleotides can be activated to form monomer-bridged-oligonucleotides that will facilitate faster extension, while the longer oligonucleotides are good templates for oligonucleotide extension. Further experiments will be required to determine the steady state distribution of VCG oligonucleotides as a function of length and sequence, over multiple cycles of replication. In addition, different rates of oligonucleotide 5′-initiation and 3′-extension could strongly influence the relative concentrations of different VCG oligonucleotides. Additional kinetic measurements will be required to determine the magnitude of such effects.

Given the observed advantage of activating the VCG oligonucleotides and the fast rate of hydrolysis of bridged N*N intermediates, some means of *in-situ* activation will clearly be required to enable continued oligonucleotide elongation and thus complete cycles of replication. Our laboratory has recently demonstrated prebiotically plausible activation and bridge-forming chemistry that allows one-pot conversion of nucleotides to bridged dinucleotides with high yield; however, this process requires repeated freeze-thaw cycles, which are known to disrupt vesicles ^23^. Therefore, a less disruptive process, compatible with vesicle integrity, may be required for VCG replication within protocells. Alternatively, if eutectic phase activation chemistry occurred in a distinct separate environment, periodic melting could potentially release fresh activated nucleotides that could flow over a population of protocells and diffuse into the vesicles while hydrolyzed nucleotides diffuse out. In such a flow system, the free 2AI generated from the formation of imidazolium-bridged species could diffuse out of the vesicles, shifting the equilibrium inside the vesicles to favor the formation of 2AI-bridged dinucleotides and monomer-bridged-oligonucleotides.

Template copying in the VCG system must proceed with sufficient fidelity to allow the inheritance of useful amounts of information; for a ribozyme on the order of 50 nucleotides in length, this implies an error rate of roughly 2% or less. Examination of the PAGE gels used to monitor primer extension reactions in our model VCG system reveals the presence of bands that do not correspond in mobility to the correct products of primer extension. In principle these bands could correspond to products of primer extension with an incorrect nucleotide, or to extension with a correct or incorrect nucleotide at the 2′-hydroxyl of the primer, or to addition of a nucleotide at the 5′-end of the primer via a 5′-5′ pyrophosphate linkage, which could be formed by attack of the 5′-phosphate of the primer on the phosphate of an activated monomer. Our experiments clearly show that 5′-5′ pyrophosphate capped oligonucleotides are generated during primer extension in the VCG system, especially from blunt ended duplexes. One of the major benefits of the VCG system is that there is no defined start or end to the genomic sequence, and oligonucleotides with a 5ʹ-capped end can still act as primers or templates. Furthermore, the synthesis of 5′-5′ pyrophosphate capped oligonucleotides suggests a straightforward way in which the evolution of ribozymes could potentiate replication. Pyrophosphate capped oligonucleotides can be substrates for ligation by ribozyme ligases, much as modern DNA and RNA ligases utilize an adenosine-5ʹ-5ʹ-pyrophosphate activated substrate.^24^ Our lab has previously evolved a ribozyme ligase that catalyzes ligation of adenylated RNAs to demonstrate the prebiotic possibility of such a mechanism.^22^

In addition to mutations induced by 3′-misincorporations, mis-priming can also be a source of mutations. A vesicle membrane that would encapsulate the VCG system and separate it from the external environment could therefore be extremely beneficial. The uptake of short oligonucleotides, such as dimers and trimers, from the external environment should not cause problems, due to competition for binding to complementary template sites. On the other hand, the uptake of longer mismatched oligonucleotides (5-8nt) could be mutagenic. This may provide a useful constraint in defining the desirable properties of protocell membranes.

Finally, we note that genome replication via the VCG model provides the raw materials necessary for spontaneous ribozyme assembly from oligonucleotides with lengths of roughly 10-20 nts. Partially overlapping pairs of such oligonucleotides can anneal with each other, after which loop-closing ligation can lead to the formation of stem-loop structures.^12^ The iteration of such processes could then lead to the assembly of complex structured RNAs, including ribozymes.

## CONCLUSION

We initially proposed the virtual circular genome (VCG) model as an approach to the nonenzymatic replication of RNA. The distributed nature of template copying in the VCG model circumvents problems associated with the replication of long linear or circular genomes. Experimental tests of the rate, extent and fidelity of template copying are clearly required to assess the viability of the VCG model. Our initial experiments show that template-directed primer extension can indeed occur within a complex synthetic VCG oligonucleotide mixture, supporting our conjecture that a fraction of annealed configurations of VCG oligonucleotides would be productive for substrate binding and reaction. The surprisingly long time course of primer extension suggests that these annealed configurations continue to rearrange spontaneously for extended times, approaching the thermodynamic minimum of full base-pairing very slowly. Our hypothesis that an exponential oligonucleotide concentration vs. length profile would support rapid replication is not supported; rather, we find that a U-shaped profile is optimal for template copying. We conclude that very short oligonucleotides must be present at high concentrations approaching their K_d_s for template binding in order to act as effective primers and helpers, while a high concentration of the longest oligonucleotides is beneficial because they are the best templates. In contrast, a high concentration of medium length oligonucleotides is counter-productive because they primarily act to occlude needed templates. We find that continued primer extension is enhanced by replenishment of hydrolyzed substrates, strongly suggesting that *in-situ* activation will be required before cycles of RNA replication can be demonstrated in a VGC system. Overall, our experiments suggest that RNA replication via the VCG model may be possible, given appropriate activation chemistry and environmental fluctuations. Additional experiments will be required to determine whether a replicating VCG system can be maintained by feeding with activated monomers, or whether an input of activated oligonucleotides is also required. We are currently exploring approaches to computational modeling of VCG replication, and to the experimental demonstration of VCG replication within model protocells.

## ACKNOWLEDGMENT

J.W.S. is an investigator of the Howard Hughes Medical Institute. This work was supported in part by grants from the Simons Foundation (290363) and the National Science Foundation (2104708) to J.W.S. The authors thank Dr. Marco Todisco for his helpful discussions and assistance regarding oligonucleotide binding affinities and melting temperatures. The authors also thank Drs. Longfei Wu and Victor S. Lelyveld for helpful comments on the manuscript.

## ABBREVIATIONS

VCG: virtual circular genome;
2-AI or *: 2-aminoimidazole or 2-aminoimidazolium;
NMR: nuclear magnetic resonance;
PAGE: polyacryla-mide gel electrophoresis;
EDC: 1-Ethyl-3-(3-dimethylaminopropyl)carbodiimide.

