## Supplemental Information for "Experimental Tests of the Virtual Circular Genome Model for Non-enzymatic RNA Replication"

**1 Materials and Methods**
**1.1 Abbreviations**
**1.2 General information**
**1.3 Selection of VCG sequence**
**1.4 Synthesis of activated nucleotides**
**1.4.1 Synthesis of 2-aminoimidazolium-bridged-dinucleotide intermediates**
**1.4.2 Synthesis of 2-aminoimidazole activated trimer**
**1.4.3 Synthesis of 2-aminoimidazole activated 3′-amino-2′,3′-dideoxyribonucleotides**
**1.5 NMR measurements of equilibration and hydrolysis of activated nucleotides**
**1.6 Radiolabeled oligonucleotides**
**1.6.1 General protocol for oligonucleotide radiolabeling and purification**
**1.6.2 5′-adenylation of radiolabeled oligonucleotides**
**1.7 Monitoring primer extension in the VCG oligonucleotide mixture**
**1.7.1 Preparation of the VCG oligo mixture**
**1.7.2 Standard VCG reactions**
**1.7.3 Primer extension reactions with preactivated VCG oligonucleotides**
**1.7.4 Phosphatase digestion**
**1.8 Melting temperature measurements**

**2 Supplementary Figures and Tables**

### 1. Materials and Methods

#### 1.1 Abbreviations

VCG, virtual circular genome
2-AI, 2-aminoimidazole
*, 2-aminoimidazole or 2-aminoimidazolium bridge
CV, column volume
HPLC, high-performance liquid chromatography
HRMS, high-resolution mass spectrometry
LC-MS, liquid chromatography-mass spectrometry
NMR, nuclear magnetic resonance
DSS, 3-(Trimethysilyl)-1-propanesulfonic acid-$d_6$ sodium salt
PAGE, polyacrylamide gel electrophoresis
Q-TOF, quadrupole time-of-flight
TEA, triethylamine
TEAB, triethylammonium bicarbonate
Tris, tris(hydroxymethyl)aminomethane
EDTA, Ethylenediaminetetraacetic acid
TBE buffer, Tris/Borate/EDTA buffer
EDC, 1-Ethyl-3-(3-dimethylaminopropyl)carbodiimide

#### 1.2 General information

All chemicals were purchased from Sigma-Aldrich (St. Louis, MO) and used without purification unless otherwise noted. All ribonucleoside 5′-monophosphates were purchased as the free-acid form from MP Biotechnology (Solon, OH). 2,2′-Dipyridyl disulfide was purchased from Combi-Blocks (San Diego, CA). 1-Hydroxy-7-azabenzotriazole was purchased from Creosalus (Louisville, KY). Adenosine triphosphate ($\gamma$-$^{32}$P labeled) was purchased from Perkin Elmer (Waltham, MA). T4 polynucleotide kinase and 5′ DNA adenylation kit were purchased from New England Biolabs (Ipswich, MA). Phosphoramidites and reagents used for solid-phase RNA synthesis were purchased from ChemGenes (Wilmington, MA) and Glen Research (Sterling, MA). Deuterated solvents were purchased from Cambridge Isotope Laboratories (Tewksbury, MA). Illustra NAP-5 Sephadex G-25 DNA grade columns were purchased from GE Healthcare (Chicago, IL)

Reverse phase flash chromatography was performed using prepacked RediSep Rf Gold C18Aq 50 g columns from Teledyne Isco (Lincoln, NE). Preparatory-scale high performance liquid chromatography (HPLC) was carried out on an Agilent 1290 HPLC system, equipped with a preparative-scale Agilent ZORBAX Eclipse-XDB C18 column (21.2x250mm, 7 μm particle size) for reversed-phase chromatography. UV melting measurements were performed on a Cary 3500 UV-Vis Spectrophotometer from Agilent. $^{1}$H, and $^{31}$P spectra were acquired on a Varian Oxford AS-400 NMR spectrometer (400 MHz for $^{1}$H, 162 MHz for $^{31}$P). Chemical shifts are reported in parts per million (ppm) values on the $\delta$ scale. $^{1}$H NMR was referenced using DSS as internal standard (0 ppm at 25 °C). All NMR spectra were recorded at 25 °C.

#### 1.3 Selection of VCG sequence

To facilitate initial testing of the VCG model, we devised a 12-nucleotide sequence containing all four canonical nucleotides using the following series of steps. Although there are $4^{12}$ (16,777,216) different 12-mers, most circular double-stranded 12-mer sequences consist of 24 related sequences, 12 from the + strand and 12 from the complementary - strand. As an example, 5′-CAAAAAAAAAAA-3′ is equivalent to twelve circularly permuted sequences such as 5′-ACAAAAAAAAAA-3′, 5′-AACAAAAAAAAA-3′, and so on, plus the 12 complementary sequences. Accounting for this 24-fold redundancy, we have a total pool of 700,274 sequences corresponding to unique non-equivalent double-stranded circular genomes.

We then applied the following filters sequentially in order to find 12-mer VCG genomes with the best chance of being able to undergo overall replication. (a) We first eliminated sequences containing the dinucleotides UU and UA in either strand, since these sequences showed very poor primer extension in our previous studies.[1] (b) We then eliminated sequences containing two or more occurrences of the pyrimidine dinucleotides CU and UC, for the same reason. (c) Next, we eliminated sequences with single nucleotide runs of length three or more, such as AAA, which could lead to mis-pairing, and (d) we eliminated sequences that exhibit a minimum free energy secondary structure with 3- or 4-nt stem-loops based on predictions using the *RNAfold* functionality within the ViennaRNA Package 2.0.[2] (e) Finally we eliminated sequences with repeated four-letter or longer words in the same strand, and repeated three-letter or longer words between the + and - strands. Removing all sequences with repeated sequences would remove all sequences, and hence, we allowed short words (≤3 nts) to be repeated in our VCG sequence.

Using the above filters, we derived 15 sequences whose GC content ranged from 58.3% to 75%. Because of its lower GC content and a distributed spread of the GC regions, we used the sequence 5′-ACACGCAUCACC-3′ for kinetic investigations into the VCG model for RNA replication in this work. A VCG sequence of size 12 can have up to 256 unique oligonucleotides with sizes varying from two to 12. Since we could not eliminate all repeats, our sequence generates 247 unique oligonucleotides ranging from 2 to 12 nucleotides in length.

#### 1.4 Synthesis of activated nucleotides

##### 1.4.1 Synthesis of 2-aminoimidazolium-bridged-dinucleotide intermediates

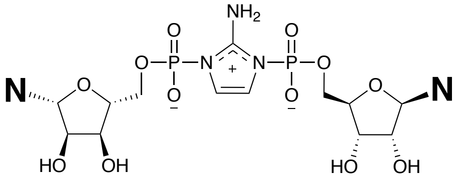


Imidazolium-bridged homo-dinucleotides (U*U, A*A, C*C, G*G) were synthesized as previously described.[1] The imidazolium-bridged dinucleotides were purified by preparative scale HPLC using a C18 reverse phase column and a solvent gradient of (A) 2 mM aqueous TEAB buffer, pH 8.0, (B) acetonitrile. The desired products were eluted between 2% and 5% B over 27 min with a flow rate of 15 mL/min. The fractions containing the product were combined, and the pH was adjusted to 8 before lyophilization.

##### 1.4.2 Synthesis of 2-aminoimidazole activated trimer

The activated trimer *GUG was synthesized as previously described.[1] The *GUG was purified by reverse-phase chromatography, with a 50g C18Aq column, over 20 CVs of 0-10% acetonitrile in 20 mM TEAB buffer (pH 8.0). The fractions containing the product were adjusted to pH 9.5~10, aliquoted into small tubes with the desired amount for one VCG reaction, and lyophilized.

##### 1.4.3 Synthesis of 2-aminoimidazole activated 3′-amino-2′,3′-dideoxyribonucleotide

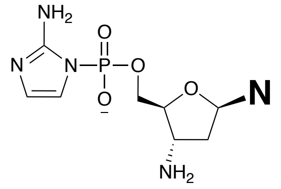


The four different monomers (3′-$NH_2$-2AIpddN) were synthesized and purified separately following the previously published protocol.[3]

#### 1.5 NMR equilibration and hydrolysis measurement

A 500 µL sample was prepared with 5 mM of each of the four bridged homo-dinucleotides and 200 mM deuterated d11-Tris-Cl (pH 8.0) in $D_2O$. $^1H$ and $^{31}P$ spectra were acquired immediately after the sample was prepared, then at 0.5, 1, 2, 4 h. Trace TEA in the bridged dinucleotides samples from purification was used as the internal standard for integration (δ 3.2 (q, $J$ = 7.3 Hz, 2H), 1.28 (t, $J$ = 7.3 Hz, 3H)). The TEA concentration was determined based on its integration ratio to the bridged dinucleotides at the initial spectra. Activated bridged dinucleotides and monomer concentrations at subsequent time points were measured by integrating corresponding $^1H$ or $^{31}P$ peaks.[1,4]

Hydrolysis under the primer extension conditions was measured with 5 mM of each of the four bridged homo-dinucleotides, 200 mM deuterated d11-Tris-Cl (pH 8.0), and 50 mM $MgCl_2$. $^{31}P$ spectra were acquired at 0.1, 0.4, 1, 2, 4, 6, 24 h. Ratios between the $^{31}P$ peaks were used to determine the concentration of each species.

#### 1.6 Radiolabeled oligonucleotides

##### 1.6.1 General protocol for radiolabeling and purification of oligonucleotides

Oligonucleotides with a 5′-OH were prepared on an Expedite 8909 DNA/RNA synthesizer and purified by reverse-phase chromatography, with a 50g C18Aq column, over 20 CVs of 1-10% acetonitrile in 20 mM TEAB buffer (pH 8.0). The synthesized oligonucleotides were 5′-labeled with $^{32}P$-$\gamma$-ATP and T4 polynucleotide kinase, followed by phenol/chloroform extraction of the kinase. Purification was performed on a Sephadex G-25 column. The labeled oligonucleotides were analyzed by 20% denaturing PAGE to ensure that all unreacted $^{32}P$- $\gamma$-ATP had been removed.

##### 1.6.2 5′-Adenylation of radiolabeled oligonucleotides

The 5′-radiolabeled oligonucleotide $^{32}$pGUGAUG was adenylated using a 5′-DNA adenylation kit (New England Biolabs) in a 20 µl volume. The solution was incubated at 65°C for 1 h before heating at 85°C for 5 min. Then the solution was purified directly on a Sephadex G-25 column and analyzed by 20% denaturing PAGE.

#### 1.7 Monitoring primer extension in the VCG oligonucleotide mixture

##### 1.7.1 Preparation of the VCG oligonucleotide mix

The 5′-phosphorylated RNA dimers, trimers, and tetramers used in this study were synthesized on a MerMade 6 DNA/RNA synthesizer (Bioautomation, Plano, TX) and purified on a 50g C18Aq column over 20 CVs of 0-10% acetonitrile in 20 mM TEAB buffer (pH 8.0). Oligonucleotides longer than 4-nts were purchased from Integrated DNA Technologies (Coralville, IA).

Stock solutions of each individual oligonucleotide were prepared first, and then stock solutions containing all oligonucleotides with the same length (from dimer to 12-mer) were prepared. The repeated dimers and trimers in the VCG sequence were prepared at a higher concentration corresponding to their multiple occurrences. Complete oligonucleotide mixtures required for a 10-µL-scale primer extension experiment were prepared from the appropriate stock solutions in a PCR tube (VWR, Radnor, PA). The solution was then lyophilized and stored at -80°C. Oligonucleotides for single template and nontemplated experiments were prepared following similar procedures using stock solutions of individual oligonucleotides.

##### 1.7.2 Standard VCG reactions

Solutions containing all 10 imidazolium-bridged dinucleotides were prepared by allowing a solution containing 10 mM of each imidazolium-bridged homo-dinucleotide(U*U, A*A, C*C, G*G) to equilibrate for 2 h on a rotator. The pre-lyophilized 10-µL-scale VCG oligo mix was dissolved in 2 µL of 1M Tris-HCl (pH 8.0), 0.5 µL of 1M $MgCl_2$, and 2.5 µL of radiolabeled oligonucleotide (<0.2 µM). The reaction was initiated by adding 5 µL of the pre-equilibrated bridged-dinucleotide mix to generate 10 µL of a solution containing the desired concentration of the VCG oligonucleotides, the radiolabeled oligonucleotide (< 0.05 µM), 50 mM $MgCl_2$, 200 mM Tris-HCl, and 20 mM total N*Ns. Immediately after the addition of the N*N mix, the reaction mixture was heated to 90°C for 10 s in a thermal cycler machine and then quickly cooled on ice for 1 min.

When the activated trimer *GUG was used, it was prepared as lyophilized powder in the desired quantity, dissolved in 5 µL pre-equilibrated 40 mM N*N, and added to the VCG mixture to initiate the reaction. Experiments with 20 mM of only one of the imidazolium-bridged homo-dinucleotides or 40 mM of 3′-$NH_2$-2AIpddN mix were initiated by adding the 2-fold concentrated substrate solution without pre-equilibration.

High temperature pulses during the course of primer extension reactions were performed following the same procedure as the initial heat pulse. To replenish mixed bridged-dinucleotides during long primer extension reactions, the desired amount of N*N stock solution was pre-equilibrated then lyophilized before being dissolved into the primer extension reaction mixture.

At each time point, a 0.5 µl aliquot of the reaction mixture was added to 25 µl of quenching buffer containing 25 mM EDTA and 1X TBE in formamide. The components of the reaction mixture were resolved by 20% (19:1) denaturing PAGE using 45×35 cm glass plates running at 50W for about 4.5 h. The gel region containing radioactive products was exposed to a phosphor-imaging plate overnight, which was then scanned using a Typhoon 9410 scanner and analyzed using ImageQuant TL software.

##### 1.7.3 Primer extension reactions with preactivated VCG oligonucleotides

The VCG oligonucleotides were first mixed in the desired ratios from the master mixes as described in 1.7.1, and then lyophilized to dryness. The powder was then re-dissolved to 20-fold of the desired final VCG concentration with 1 equivalent of 2-aminoimidazole (pH 7) and 20 equivalents of EDC (pH 7). The resultant mixture was fast-frozen with liquid nitrogen and then incubated at -15°C overnight for eutectic phase concentration and activation. On the next day, the VCG reaction was performed similarly to section 1.7.2, except that 0.5 μL of the pre-activated 20-fold VCG mixture was mixed with 2 μL of 1M Tris-HCl (pH 8.0), 0.5 μL of 1M $MgCl_2$, and 2 μL of radiolabeled oligonucleotide. Then 5 μL of pre-equilibrated 40 mM N*N was added, followed by a heat pulse to initiate the reaction. Time points were collected and analyzed as described in 1.7.2.

##### 1.7.4 Phosphatase digestion

To identify 5′-modified oligonucleotides, unmodified 5′-phosphates were removed by phosphatase treatment as follows. All VCG and the nontemplated extension reactions were performed at 10 μL scale as indicated above. At 24 hours, the sample were mixed with 40 μL ethanol and 2 μL of 0.5 M EDTA in a siliconized microcentrifuge vial. The vial was cooled on dry ice for 2 hours, after which it was centrifuged at 21,000 g for 10 minutes. The resulting pellet was washed twice with 80% ethanol, and then dried on SpeedVac vacuum concentrator for 10 minutes. The pellet was resolubilized in 10 μL water. A 0.5 μL aliquot was treated with the Quick CIP (New England Biolabs) phosphatase in 20 μL. After heat deactivation of the phosphatase, the mixture was added to 20 μL buffer containing 8M Urea, 1X TBE, and 25 mM EDTA. The standard VCG and nontemplated extension reactions were also quenched with the same buffer and diluted to the same concentration as the phosphatase digested samples. The samples were resolved next to each other on a 20% (19:1) denaturing PAGE with 45×35 cm glass plates. The gel region containing radioactive samples was cut, wrapped between Saran Wrap sheets, and exposed to a phosphor imaging plate overnight. The gel was scanned using a Typhoon 9410 scanner and analyzed using ImageQuant TL software.

#### 1.8 Melting temperature measurements

Melting temperatures were measured using an Agilent 3500 UV-Vis Spectrophotometer. For each pair of complementary oligonucleotides, samples were prepared with 1 μM of the target oligonucleotide and 1 μM of its complementary strand in 50 mM $MgCl_2$ and 200 mM Tris-HCl (pH 8.0). Melting curves were collected by following absorbance at 260 nm as a function of temperature using a temperature ramp of 3°C/min. The readings were collected in heating-cooling cycles with respect to a control sample containing 50 mM $MgCl_2$ and 200 mM Tris-HCl (pH 8.0). The melting temperatures were calculated from a double baseline, two-state fit of the raw melt data, and then averaged from at least four replicates at the same condition.

#### 2 Supplementary Figures and Tables

**Figure S1.**

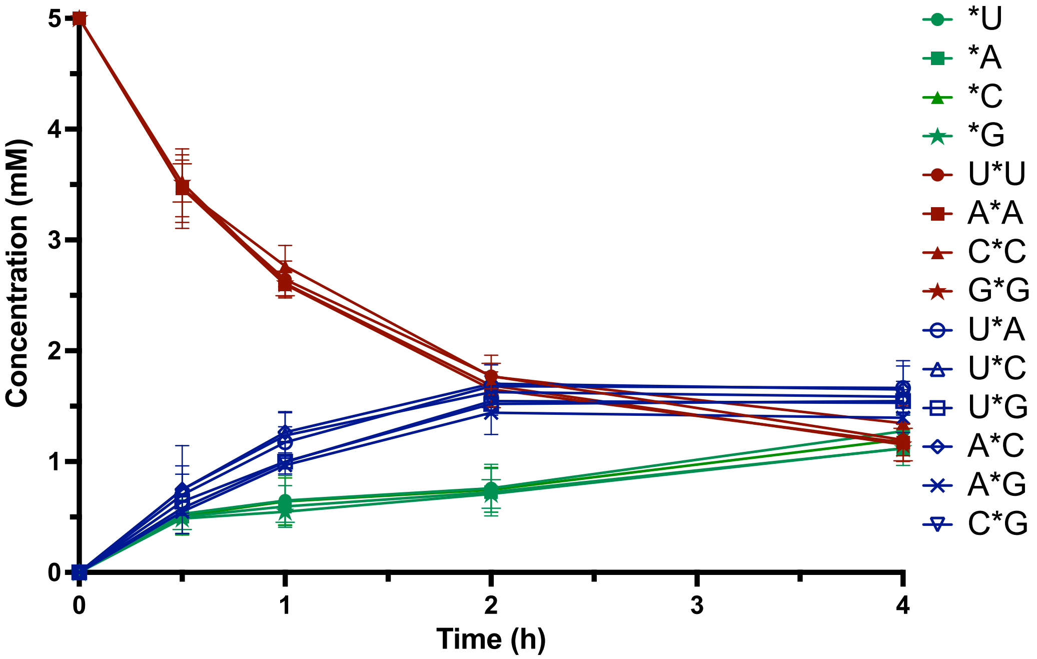


**Figure S1.** Equilibration of four bridged homo-dinucleotides (U*U, A*A, C*C, G*G) determined by NMR. Experiments were started with 5 mM of each bridged homo-dinucleotides and 200 mM d11-Tris-DCl (pH 8.0) in $D_2O$. DSS (δ 3.2 (s, $J$ = 0 Hz)) was used as the internal standard for chemical shifts. Trace TEA in the bridged dinucleotides samples from purification was used as the internal standard for integration at different time points (δ 3.2 (q, $J$ = 7.3 Hz, 2H), 1.28 (t, $J$ = 7.3 Hz, 3H)). Error bars were derived from triplicate experiments.

**Figure S2.**

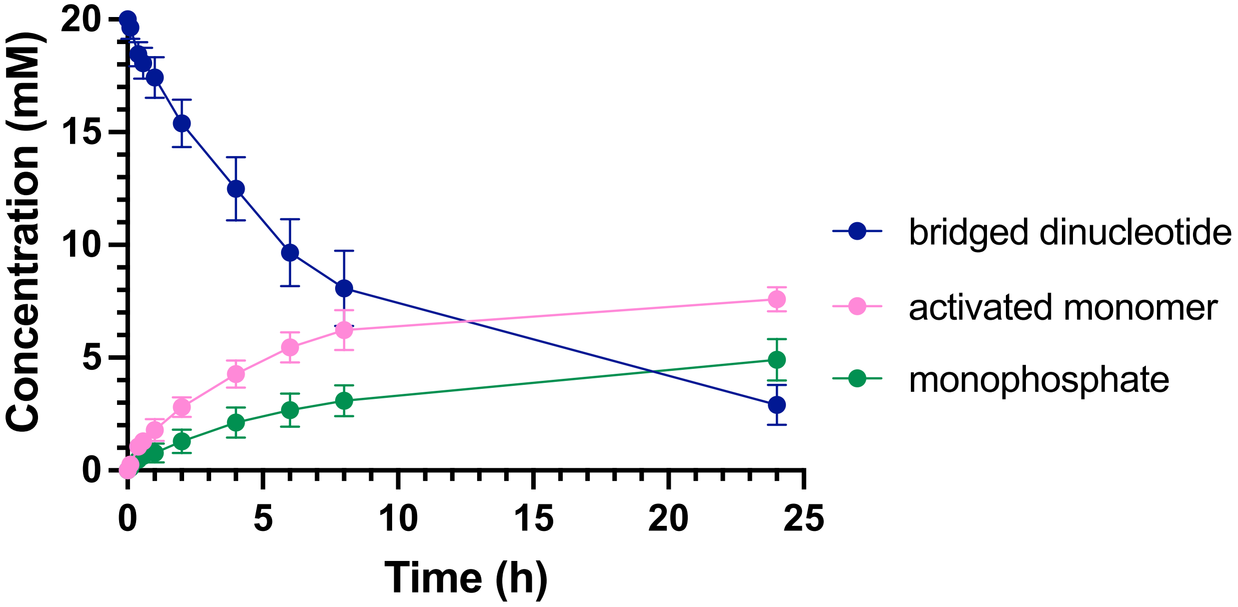


**Figure S2.** Hydrolysis of bridged intermediates in VCG primer extension conditions determined by $^{31}P$ NMR. Experiments were started with 5 mM of each four bridged homo-dinucleotides (U*U, A*A, C*C, G*G), 50 mM $MgCl_2$, and 200 mM d11-Tris-DCl (pH 8.0) in $D_2O$. The sum of concentrations of all of the bridged dinucleotides, all of the activated monomers, and all of the monophosphates are plotted. Error bars were derived from triplicate experiments.

**Figure S3.**

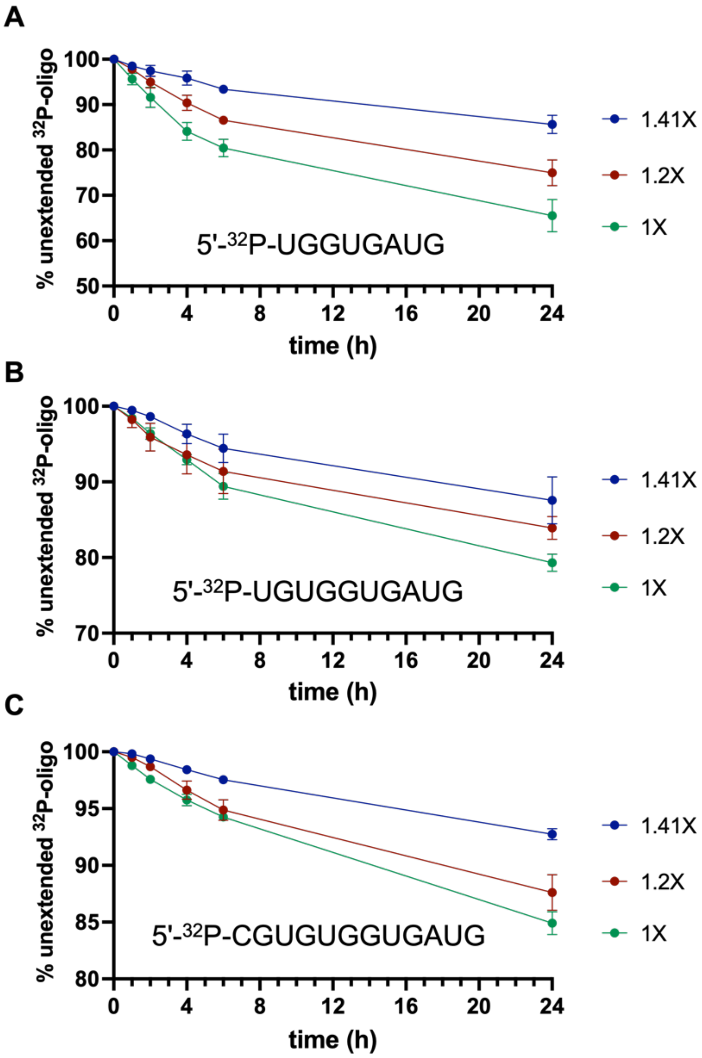


**Figure S3.** Primer extension of oligonucleotides of different lengths (A) 8mer (B) 10mer (C) 12mer in VCG mixes with different concentration vs. length gradients. The extent of extension in the VCG mix was measured by doping in a $^{32}P$ labeled oligonucleotide, whose sequence was labeled on the figure. All reactions were conducted at room temperature, with 50 mM MgCl2, 200 mM Tris-HCl (pH 8.0), and 20 mM pre-equilibrated N*N. See Table S1 for detailed oligonucleotide concentrations in the different VCG mixes.

**Figure S4.**

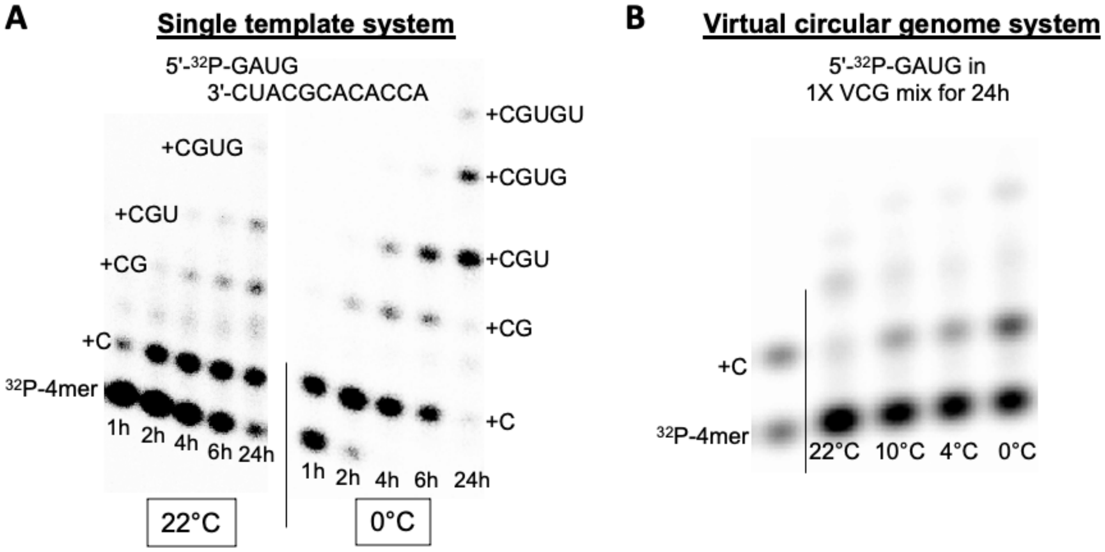


**Figure S4.** Primer extension of a labeled tetranucleotide, on a single template and in the VCG system. (**A**) Positive control of 5′-$^{32}$P-GAUG extension on a single template at 22°C and 0°C. The reaction contained 1μM of pACCACACGCAUC (v1202) template and 1 μM of pGAUG (v413) primer, doped with 5′-$^{32}$P-GAUG. (**B**) Primer extension of 5′-$^{32}$P-GAUG for 24h in the 1X VCG oligonucleotide mix at different temperatures. Authentic standards were run in the left lane for comparison. All reactions were conducted at 50 mM $MgCl_2$, 200 mM Tris-HCl (pH 8.0), and with 20 mM pre-equilibrated N*N.

**Figure S5.**

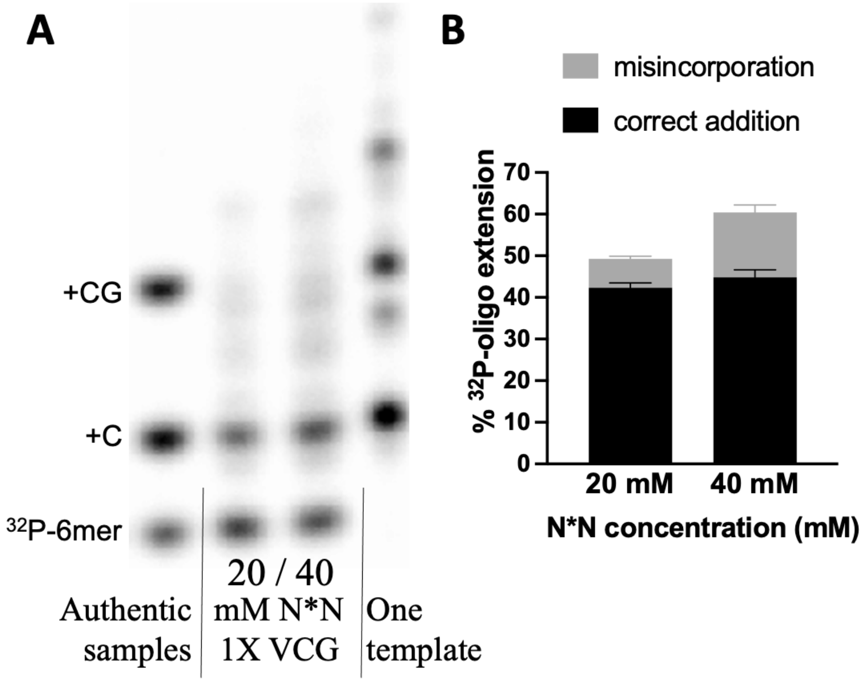


**Figure S5.** A higher concentration of activated bridged dinucleotides induces a higher level of misincorporation. (a) PAGE analysis of 5′-$^{32}$P-GUGAUG extension in 1X VCG mixture with 20 and 40 mM total N*Ns. Correct incorporations were identified by comparison with radiolabeled authentic samples and a single-template positive control. The reactions were incubated for 24 h. (b) Bar graph illustrating the percentage of correct extension and misincorporation.

**Figure S6.**

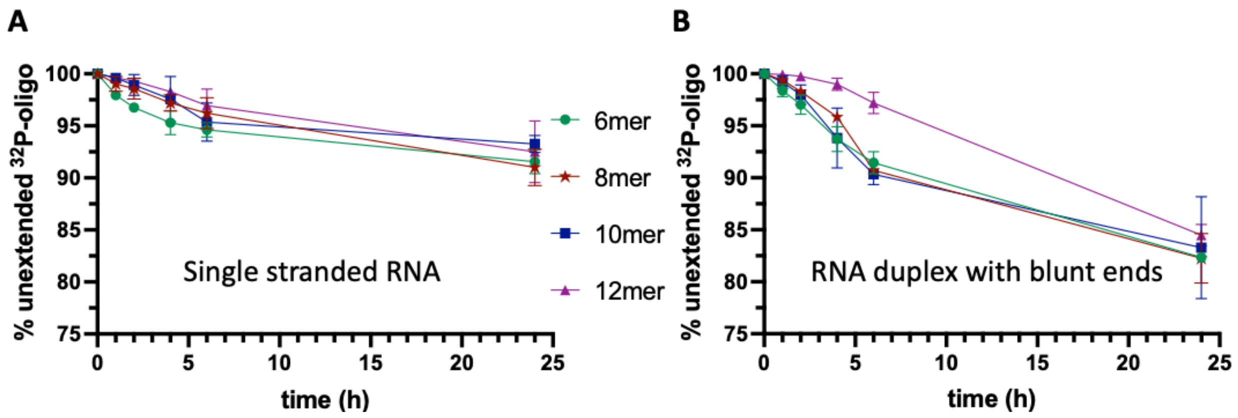


**Figure S6**. Nontemplated reactions performed with equilibrated mixture of imidazolium-bridged dinucleotides. (A) Nontemplated primer extension measured with individual single stranded oligonucleotides. (B) Nontemplated primer extension measured in the presence of the complementary oligonucleotide. The sequences of the $^{32}$P-labelled oligonucleotides are listed in **Figure 4**. All reactions were conducted for 24 h at room temperature, in 50 mM $MgCl_2$, 200 mM Tris-Cl (pH 8.0), and 20 mM total N*Ns.

**Figure S7.**

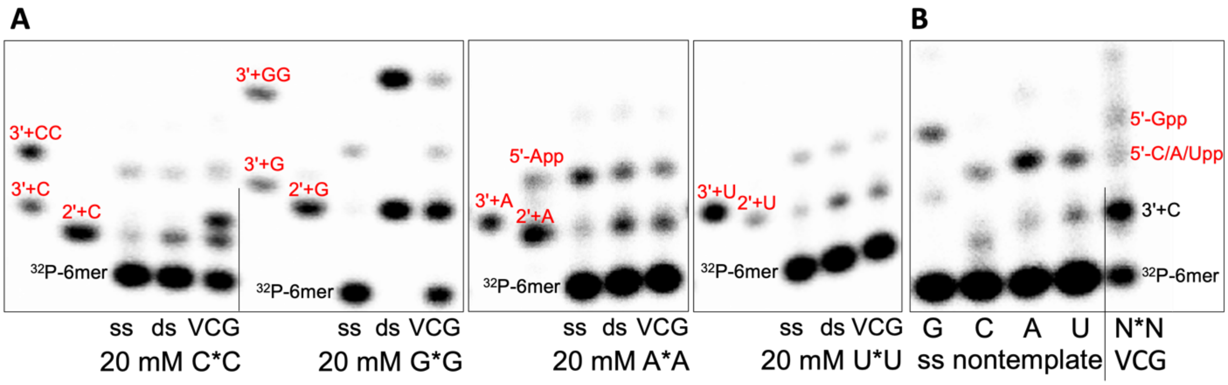


**Figure S7.** Characterization of nontemplated primer extension and misincorporations in virtual circular genome system. (A) PAGE analysis of 5′-$^{32}$P-GUGAUG reacted in solution as an isolated single stranded oligonucleotide (ss), in the presence of the complementary strand (ds), or in the presence of the 1X VCG oligonucleotide mixture, with 20 mM of the indicated imidazolium-bridged homo-dinucleotides. Authentic standards in the left two lanes are labelled in red to help identify the 5′-$^{32}$P-GUGAUG primer-extension product. (B) Aliquots of the four ss nontemplated reactions were run next to an aliquot of a standard 1X VCG primer-extension reaction. The top band of the ss nontemplated extension is the 5′-pyrophosphate (identified on the basis of resistance to phosphatase digestion), while the middle band is the product of 2′-extension with the indicated nucleotide. The ss nontemplated reactions were performed with each of the bridged homo-dinucleotides, while the VCG primer-extension reaction was with the equilibrated mixture of imidazolium-bridged dinucleotides. All reactions were conducted for 24h at room temperature, in 50 mM $MgCl_2$, 200 mM Tris-Cl (pH 8.0), and 20 mM of bridged dinucleotides.

**Figure S8.**

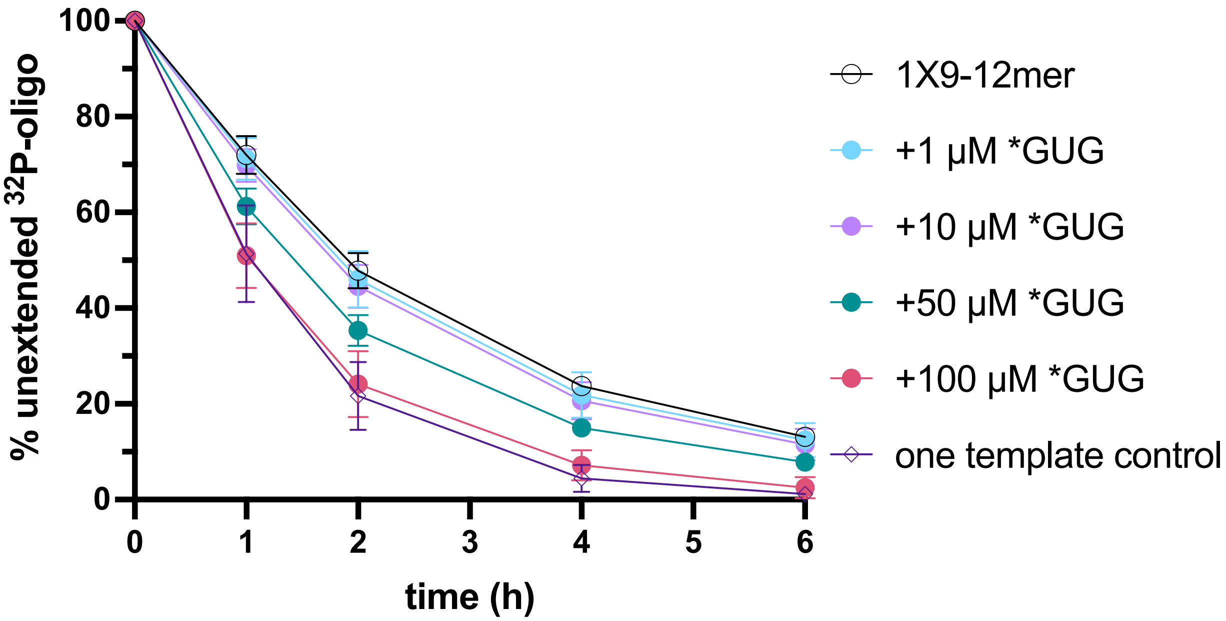


**Figure S8.** The extension of a 5′-$^{32}$P-GUGAUG inside the 1X9-12mer partial VCG system with different concentrations of *GUG and 20 mM of pre-equilibrated N*N. The extension of the one-template positive control with 20 mM N*Ns (as in Figure 2A) was also plotted for comparison. All reactions were conducted at room temperature, in 50 mM $MgCl_2$ and 200 mM Tris-HCl (pH 8.0).

**Figure S9.**

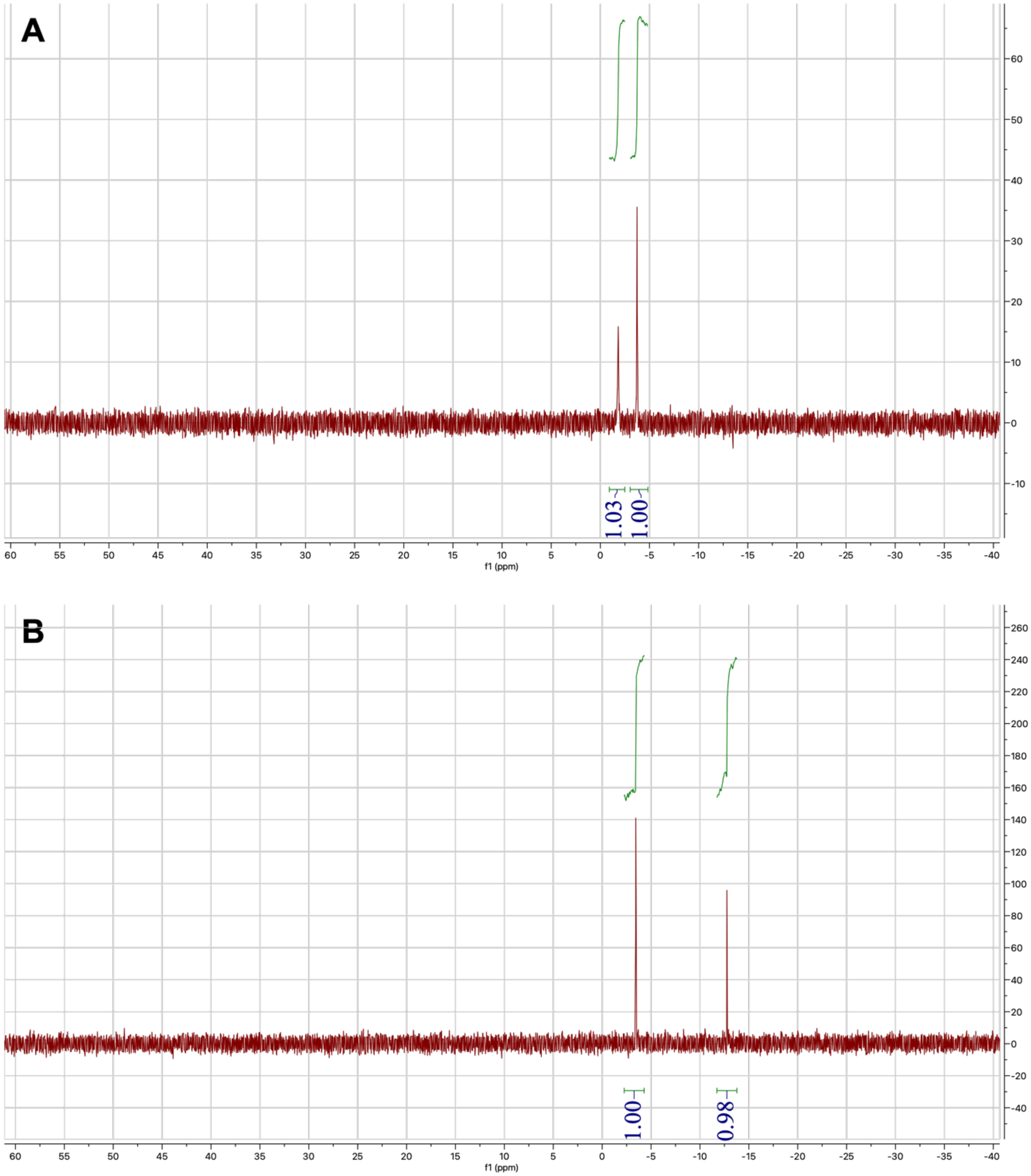


**Figure S9.** $^{32}P$ NMR of overnight eutectic phase EDC activated pCG. (A) Pure pCG before the reaction. (B) Activated pCG after the overnight EDC activation. The peak at -1.81 ppm corresponds to the 5′-phosphate of pCG while the peak at -12.76 ppm corresponds to the 5′-phosphate activated by 2AI. The peak at -3.44 ppm corresponds to the internal phosphate between the C and G nucleotides. The reaction was performed with 60 mM pCG, 1 equiv. of 2AI, and 20 equiv. of EDC at pH 7, following the same procedure for VCG activation described in 1.7.3.

**Table S1. Sequences of all RNA oligonucleotides in the VCG system used in this study.**

| | | | | | | | | | |
|---|---|---|---|---|---|---|---|---|---|
| v201 | pAC | v301 | pACA | v401 | pACAC | v501 | pACACG | v601 | pACACGC |
| v202 | pAU | v302 | pACC | v402 | pACCA | v502 | pACCAC | v602 | pACCACA |
| v203 | pCA | v303 | pACG | v403 | pACGC | v503 | pACGCA | v603 | pACGCAU |
| v204 | pCC | v304 | pAUC | v404 | pAUCA | v504 | pAUCAC | v604 | pAUCACC |
| v205 | pCG | v305 | pAUG | v405 | pAUGC | v505 | pAUGCG | v605 | pAUGCGU |
| v206 | pGA | v306 | pCAC | v406 | pCACA | v506 | pCACAC | v606 | pCACACG |
| v207 | pGC | v307 | pCAU | v407 | pCACC | v507 | pCACCA | v607 | pCACCAC |
| v208 | pGG | v308 | pCCA | v408 | pCACG | v508 | pCACGC | v608 | pCACGCA |
| v209 | pGU | v309 | pCGC | v409 | pCAUC | v509 | pCAUCA | v609 | pCAUCAC |
| v210 | pUC | v310 | pCGU | v410 | pCCAC | v510 | pCCACA | v610 | pCCACAC |
| v211 | pUG | v311 | pGAU | v411 | pCGCA | v511 | pCGCAU | v611 | pCGCAUC |
| | | v312 | pGCA | v412 | pCGUG | v512 | pCGUGU | v612 | pCGUGUG |
| | | v313 | pGCG | v413 | pGAUG | v513 | pGAUGC | v613 | pGAUGCG |
| | | v314 | pGGU | v414 | pGCAU | v514 | pGCAUC | v614 | pGCAUCA |
| | | v315 | pGUG | v415 | pGCGU | v515 | pGCGUG | v615 | pGCGUGU |
| | | v316 | pUCA | v416 | pGGUG | v516 | pGGUGA | v616 | pGGUGAU |
| | | v317 | pUGA | v417 | pGUGA | v517 | pGUGAU | v617 | pGUGAUG |
| | | v318 | pUGC | v418 | pGUGG | v518 | pGUGGU | v618 | pGUGGUG |
| | | v319 | pUGG | v419 | pGUGU | v519 | pGUGUG | v619 | pGUGUGG |
| | | v320 | pUGU | v420 | pUCAC | v520 | pUCACC | v620 | pUCACCA |
| | | | | v421 | pUGAU | v521 | pUGAUG | v621 | pUGAUGC |
| | | | | v422 | pUGCG | v522 | pUGCGU | v622 | pUGCGUG |
| | | | | v423 | pUGGU | v523 | pUGGUG | v623 | pUGGUGA |
| | | | | v424 | pUGUG | v524 | pUGUGG | v624 | pUGUGGU |

**Table S1 Continued.**

| | | | | | |
|---|---|---|---|---|---|
| v701 | pACACGCA | v801 | pACACGCAU | v901 | pACACGCAUC |
| v702 | pACCACAC | v802 | pACCACACG | v902 | pACCACACGC |
| v703 | pACGCAUC | v803 | pACGCAUCA | v903 | pACGCAUCAC |
| v704 | pAUCACCA | v804 | pAUCACCAC | v904 | pAUCACCACA |
| v705 | pAUGCGUG | v805 | pAUGCGUGU | v905 | pAUGCGUGUG |
| v706 | pCACACGC | v806 | pCACACGCA | v906 | pCACACGCAU |
| v707 | pCACCACA | v807 | pCACCACAC | v907 | pCACCACACG |
| v708 | pCACGCAU | v808 | pCACGCAUC | v908 | pCACGCAUCA |
| v709 | pCAUCACC | v809 | pCAUCACCA | v909 | pCAUCACCAC |
| v710 | pCCACACG | v810 | pCCACACGC | v910 | pCCACACGCA |
| v711 | pCGCAUCA | v811 | pCGCAUCAC | v911 | pCGCAUCACC |
| v712 | pCGUGUGG | v812 | pCGUGUGGU | v912 | pCGUGUGGUG |
| v713 | pGAUGCGU | v813 | pGAUGCGUG | v913 | pGAUGCGUGU |
| v714 | pGCAUCAC | v814 | pGCAUCACC | v914 | pGCAUCACCA |
| v715 | pGCGUGUG | v815 | pGCGUGUGG | v915 | pGCGUGUGGU |
| v716 | pGGUGAUG | v816 | pGGUGAUGC | v916 | pGGUGAUGCG |
| v717 | pGUGAUGC | v817 | pGUGAUGCG | v917 | pGUGAUGCGU |
| v718 | pGUGGUGA | v818 | pGUGGUGAU | v918 | pGUGGUGAUG |
| v719 | pGUGUGGU | v819 | pGUGUGGUG | v919 | pGUGUGGUGA |
| v720 | pUCACCAC | v820 | pUCACCACA | v920 | pUCACCACAC |
| v721 | pUGAUGCG | v821 | pUGAUGCGU | v921 | pUGAUGCGUG |
| v722 | pUGCGUGU | v822 | pUGCGUGUG | v922 | pUGCGUGUGG |
| v723 | pUGGUGAU | v823 | pUGGUGAUG | v923 | pUGGUGAUGC |
| v724 | pUGUGGUG | v824 | pUGUGGUGA | v924 | pUGUGGUGAU |

**Table S1 Continued.**

| | | | | | |
|---|---|---|---|---|---|
| v1001 | pACACGCAUCA | v1101 | pACACGCAUCAC | v1201 | pACACGCAUCACC |
| v1002 | pACCACACGCA | v1102 | pACCACACGCAU | v1202 | pACCACACGCAUC |
| v1003 | pACGCAUCACC | v1103 | pACGCAUCACCA | v1203 | pACGCAUCACCAC |
| v1004 | pAUCACCACAC | v1104 | pAUCACCACACG | v1204 | pAUCACCACACGC |
| v1005 | pAUGCGUGUGG | v1105 | pAUGCGUGUGGU | v1205 | pAUGCGUGUGGUG |
| v1006 | pCACACGCAUC | v1106 | pCACACGCAUCA | v1206 | pCACACGCAUCAC |
| v1007 | pCACCACACGC | v1107 | pCACCACACGCA | v1207 | pCACCACACGCAU |
| v1008 | pCACGCAUCAC | v1108 | pCACGCAUCACC | v1208 | pCACGCAUCACCA |
| v1009 | pCAUCACCACA | v1109 | pCAUCACCACAC | v1209 | pCAUCACCACACG |
| v1010 | pCCACACGCAU | v1110 | pCCACACGCAUC | v1210 | pCCACACGCAUCA |
| v1011 | pCGCAUCACCA | v1111 | pCGCAUCACCAC | v1211 | pCGCAUCACCACA |
| v1012 | pCGUGUGGUGA | v1112 | pCGUGUGGUGAU | v1212 | pCGUGUGGUGAUG |
| v1013 | pGAUGCGUGUG | v1113 | pGAUGCGUGUGG | v1213 | pGAUGCGUGUGGU |
| v1014 | pGCAUCACCAC | v1114 | pGCAUCACCACA | v1214 | pGCAUCACCACAC |
| v1015 | pGCGUGUGGUG | v1115 | pGCGUGUGGUGA | v1215 | pGCGUGUGGUGAU |
| v1016 | pGGUGAUGCGU | v1116 | pGGUGAUGCGUG | v1216 | pGGUGAUGCGUGU |
| v1017 | pGUGAUGCGUG | v1117 | pGUGAUGCGUGU | v1217 | pGUGAUGCGUGUG |
| v1018 | pGUGGUGAUGC | v1118 | pGUGGUGAUGCG | v1218 | pGUGGUGAUGCGU |
| v1019 | pGUGUGGUGAU | v1119 | pGUGUGGUGAUG | v1219 | pGUGUGGUGAUGC |
| v1020 | pUCACCACACG | v1120 | pUCACCACACGC | v1220 | pUCACCACACGCA |
| v1021 | pUGAUGCGUGU | v1121 | pUGAUGCGUGUG | v1221 | pUGAUGCGUGUGG |
| v1022 | pUGCGUGUGGU | v1122 | pUGCGUGUGGUG | v1222 | pUGCGUGUGGUGA |
| v1023 | pUGGUGAUGCG | v1123 | pUGGUGAUGCGU | v1223 | pUGGUGAUGCGUG |
| v1024 | pUGUGGUGAUG | v1124 | pUGUGGUGAUGC | v1224 | pUGUGGUGAUGCG |

**Table S2. Concentrations of oligonucleotides of each length in different VCG oligo mixes.**

| | 1X | 1.2 X | 1.41 X | 0.83 X | 3-fold 1X | 1/3-fold 1X | 2-8mer | 9-12mer | U-shape |
|---|---|---|---|---|---|---|---|---|---|
| Component | Concentration (μM) | | | | | | | | |
| pNpN | 1 | 6.19 | 32 | 0.16 | 3 | 0.33 | 1 | - | 4 |
| pNpNpN | 1 | 5.16 | 22.63 | 0.19 | 3 | 0.33 | 1 | - | 2.83 |
| (pN)4 | 1 | 4.3 | 16 | 0.23 | 3 | 0.33 | 1 | - | 2 |
| (pN)5 | 1 | 3.58 | 11.31 | 0.28 | 3 | 0.33 | 1 | - | 1.41 |
| (pN)6 | 1 | 2.99 | 8 | 0.34 | 3 | 0.33 | 1 | - | 1 |
| (pN)7 | 1 | 2.49 | 5.66 | 0.40 | 3 | 0.33 | 1 | - | 1.41 |
| (pN)8 | 1 | 2.07 | 4 | 0.48 | 3 | 0.33 | 1 | - | 2 |
| (pN)9 | 1 | 1.73 | 2.83 | 0.58 | 3 | 0.33 | - | 1 | 2.83 |
| (pN)10 | 1 | 1.44 | 2 | 0.69 | 3 | 0.33 | - | 1 | 4 |
| (pN)11 | 1 | 1.2 | 1.41 | 0.83 | 3 | 0.33 | - | 1 | 5.66 |
| (pN)12 | 1 | 1 | 1 | 1 | 3 | 0.33 | - | 1 | 8 |
